# The piRNA pathway mediates transcriptional silencing of LTR retrotransposons in ovaries and somatic tissues of *Aedes* mosquitoes

**DOI:** 10.1101/2024.11.28.623418

**Authors:** Ezgi Taşköprü, Nynke B. van Eijk, Femke A. H. van Hout, Marianna Bacchi, Gijs J. Overheul, Charlotte Linthout, Constantianus J. M. Koenraadt, Jieqiong Qu, Pascal Miesen, Rebecca Halbach, Ronald P. van Rij

## Abstract

The PIWI-interacting RNA (piRNA) pathway preserves genomic integrity by suppressing transposable elements in animal germlines. Despite its well-established function in the animal germline, piRNAs and PIWI proteins are expressed in somatic tissues across arthropod species, and their functions outside the gonads remain poorly understood. *Aedes albopictus* mosquitoes express four PIWI genes, *Piwi4*, *Piwi5*, *Piwi6* and *Ago3,* in both gonadal and somatic tissues. Here, we generated *Piwi6* knockout *Ae. albopictus* cell lines and observed a substantial upregulation of Long Terminal Repeat (LTR)-retrotransposons, including a full-length endogenous retrovirus that we named *AalERV1*. Nascent RNA-sequencing and CUT&Tag analyses revealed that Piwi6 silences *AalERV1* transcriptionally by guiding the deposition of the repressive H3K9me3 histone mark. Consistently, Piwi6 localized to both the cytoplasm and nucleus, with sequences in the intrinsically disordered region guiding nuclear translocation. Reintroduction of full-length GFP-Piwi6, but not a mutant GFP-Piwi6 defective in nuclear localization, rescued *AalERV1* derepression in *Piwi6* knockout cells. Importantly, the control of *AalERV1* was recapitulated *in vivo* as *Piwi6* knockdown increased *AalERV1* expression in both ovaries and somatic tissues of *Ae. albopictus* mosquitoes. These results establish *Aedes* mosquitoes as a model to study nuclear PIWI functions and suggest that somatic piRNA-mediated transposon silencing is evolutionarily conserved across arthropod species.

## Introduction

Transposable elements (TEs) are selfish genetic elements capable of moving within host genomes, integrating into new sites and causing mutations that threaten the cell’s genetic identity. In most animal species, the PIWI-interacting RNA (piRNA) pathway serves as a specialized defense mechanism to control TEs in the gonads. The piRNA pathway has been extensively studied in the germline of *Drosophila melanogaster*, where 24–32 nt sized piRNAs originate from discrete genomic regions known as piRNA clusters [1–3]. These piRNA clusters are rich in repetitive sequences derived from past transposon invasions and produce piRNAs that associate with PIWI proteins and guide them to their targets through sequence complementarity [4,5].

Target engaged PIWI-piRNA complexes can induce silencing via two distinct mechanisms: post-transcriptional gene silencing by degradation of existing RNAs and transcriptional gene silencing by establishing repressive chromatin at transposon loci. In *Drosophila*, these modes of silencing are executed by distinct PIWI proteins. Post-transcriptional gene silencing is mediated by Aubergine (Aub) and Argonaute3 (Ago3), which cleave transposon transcripts and participate in a piRNA amplification loop termed the ping-pong cycle [6]. This process occurs in the cytoplasm of germ cells in a membraneless organelle called the nuage [7]. In contrast, transcriptional gene silencing is mediated by Piwi and occurs in the nucleus in a slicing-independent manner. Once loaded with a piRNA, Piwi translocates to the nucleus where it recognizes nascent transposon transcripts and recruits chromatin modifiers, resulting in heterochromatin formation [8–12].

In contrast to *Drosophila melanogaster*, the function of the piRNA pathway in most arthropod species extends beyond the gonads [13]. For example, multiple PIWI genes in *Aedes* mosquitoes are expressed both in somatic tissues and the gonads [14,15]. Interestingly, seven and nine PIWI genes are annotated in the current genome annotations of the closely related *Aedes aegypti* (yellow fever mosquito) and *Aedes albopictus* (Asian tiger mosquito), respectively [14–18], compared to the three PIWI genes of *Drosophila.* This gene expansion suggests potential functional diversification of the piRNA pathway in *Aedes* mosquitoes. Indeed, PIWI proteins associate with piRNAs from diverse sources, including cytoplasmic viral RNA, non-retroviral endogenous viral elements (nrEVEs), protein-coding genes, and a highly conserved satellite repeat, in addition to TEs [19–23].

In *Aedes* mosquitoes, the PIWI genes *Ago3*, *Piwi4*, *Piwi5* and *Piwi6* are ubiquitously expressed both in gonads and non-gonadal tissues [14,15]. Whereas functions of Ago3, Piwi4 and Piwi5 have been characterized in *Ae. aegypti* [19,20,24,25], the function of Piwi6 has thus far remained unclear. We previously showed that *Ae. aegypti* Piwi6 associates with TE derived piRNAs, an unexpected finding given that *Piwi6* knockdown did not affect piRNA biogenesis. In contrast, Piwi5 and Ago3 were found to mediate TE derived piRNA biogenesis and, accordingly, to associate with these piRNAs, with Ago3 exclusively required for production of piRNAs by ping-pong amplification [25]. The role of Piwi6 in *de novo* production of piRNAs from viral RNA seems to be minor and virus dependent. For example, *Piwi6* knockdown did not affect viral piRNA biogenesis in infections with Sindbis virus and Semliki Forest virus [25,26], while it reduced dengue virus-derived piRNA levels, albeit to a lesser extent than *Piwi5* and *Ago3* knockdown [27]. Furthermore, mass spectrometric analyses indicated that Piwi6 interacts with piRNA biogenesis factors such Yb, vreteno, minotaur, tejas, as did Piwi4, Piwi5 and Ago3. Intriguingly however, Piwi6 was the only protein that also interacted with proteins with putative nuclear functions in chromatin assembly, nucleic acid binding, or regulation of transcription by RNA polymerase II [28], suggesting that Piwi6 mediates transcriptional gene silencing.

In this study, we investigated Piwi6 function in *Ae. albopictus* mosquitoes using CRISPR-Cas9 mediated knockouts (KO) in U4.4 cells, a cell line that mimics PIWI protein and piRNA expression of somatic tissues [29]. Through small RNA and mRNA sequencing, nascent RNA-sequencing and CUT&Tag, we found that Piwi6 mediates transcriptional gene silencing of LTR retrotransposons, including a full-length LTR/Gypsy-like endogenous retrovirus that we named *AalERV1*. Furthermore, structure-guided mutational analyses revealed that the intrinsically disordered region (IDR) of Piwi6 is crucial for nuclear localization and transcriptional gene silencing. Our results establish that the *Aedes* piRNA pathway has a nuclear arm that functions in transcriptional silencing of TEs in the soma, which evolved independently of the nuclear function of *Drosophila* Piwi.

## Materials and Methods

### Cells and viruses

*Ae. albopictus* U4.4 cells (RRID:CVCL_Z820) were cultured at 28 °C in Leibovitz’s L-15 medium (Gibco) supplemented with 10% heat-inactivated fetal bovine serum (FBS; Gibco), 2% tryptose phosphate broth solution (Sigma-Aldrich), 1x MEM non-essential amino acids (Gibco), and 50 U/mL penicillin and 50 μg/mL streptomycin (pen/strep, Gibco). The cell lines were tested regularly for mycoplasma contamination. When indicated, wild-type, control and *Piwi6* KO cells were seeded in 24-well plates and infected with recombinant Sindbis virus expressing GFP from a duplicated subgenomic promoter (SINV-GFP, produced from plasmid pTE-3’2J-GFP [30]) at a multiplicity of infection (MOI) of 1 for 48 h.

### Generation of *Piwi6* knockout cells

U4.4 control and *Piwi6* knockout (KO) cells were obtained using CRISPR/Cas9 as described previously [31]. Briefly, single guide RNA sequences (Supplementary Table S1) targeting *Ae. albopictus Piwi6* (*AALFPA_065618*) were cloned into the pAc-Cas9-AalbU6.2 plasmid described in [31], replacing the 5’-GGAAGAGCGAGCTCTTCC-3’ sequence that was used as negative control. The resulting plasmids were transfected into WT cells, and 2 days later, transfected cells were selected by 20 μg/mL puromycin treatment. The editing efficiency of specific guides was assessed on agarose gel after PCR on genomic DNA. The cells transfected with the guide sequences that showed the highest editing efficiency were seeded at a density of a single cell per well in the absence of puromycin. After 3 weeks, single-cell clones containing insertions/deletions in the targeted *Piwi6* locus were identified by PCR and Sanger sequencing, resulting in two clones containing out-of-frame mutations: *Piwi6* KO#1 (guide 1 - clone #10) and *Piwi6* KO#2 (guide 1 - clone #52). Control clones, Ctrl #1 (clone #1) and Ctrl #2 (clone #6), were generated in parallel with the *Piwi6* KO cells, by transfecting U4.4 cells with pAc-Cas9-AalbU6.2 plasmid lacking sgRNA sequences.

### Plasmids

Plasmids encoding GFP-tagged wild-type *Piwi6* were generated by inserting the cDNA encoding for *Ae. albopictus Piwi6* gene into the pUb-GW-GFP plasmid [32]. PCR was used to generate the backbone and insert the *Piwi6* sequence using CloneAmp HiFi PCR premix (Takara Bio) and the primers listed in Supplementary Table S2.

GFP-tagged *Piwi6* deletion constructs (IDR, NLS mutants, sequential deletions in the IDR) and mutant *Piwi6* (AS* and YK*) were generated by site-directed mutagenesis using the In-Fusion HD cloning kit (Takara Bio) with primers listed in Supplementary Table S2. Sequences of the constructs were confirmed by Sanger sequencing and protein expression was verified by western blotting.

### RNA isolation

Cells were harvested in 1 mL RNA-Solv Reagent (Omega Bio-tek). To initiate phase separation, 200 μL chloroform was added and the samples were mixed vigorously with a vortex. The samples were centrifuged at 18,000 x *g* for 20 min, and RNA was precipitated from the aqueous phase using ice-cold isopropanol with 20 μg glycogen (Invitrogen) by incubation on ice for 30 min to 1 h. RNA was precipitated by centrifugation at 18,000 x *g* for 30 min at 4°C. The RNA pellets were washed with 85% ethanol, air-dried, and dissolved in nuclease-free water.

### Reverse transcription and quantitative PCR

cDNA was produced from 1 μg of total RNA, treated with DNaseI (Ambion) according to the manufacturer’s instructions, and reverse transcribed using the TaqMan MultiScribe Reverse Transcription Kit (Applied Biosystems) with oligo(dT) primers. The cDNA was used as a template in both qPCR and end-point PCR reactions. qPCR was performed on a LightCycler 480 (Roche) using the GoTaq qPCR Mix (Promega). Target gene expression was normalized to an internal control, the *RPL5* gene, and the fold changes compared to controls were calculated using the 2^(-ΔΔCT)^ method [33]. The primers used in qPCR reactions are listed in Supplementary Table S3.

To analyze *PIWI* gene expression in mosquito tissues, 1 μg of total RNA was reverse transcribed as described above, and PCR was performed using GO-Taq Flexi DNA Polymerase (Promega) and the primers listed in Supplementary Table S5. A reaction in the absence of the reverse transcriptase enzyme was included as a negative control to detect genomic DNA contamination.

To measure SINV RNA copies in WT, Ctrl and *Piwi6* KO cells, total RNA was reverse transcribed as described above and qPCR was performed on a LightCycler 480 (Roche) using the GoTaq qPCR Mix (Promega). Relative SINV RNA copies were calculated by using a standard curve generated by performing dilution series of plasmid containing SINV *NSP4* sequence.

### Actinomycin Treatment

WT, *Piwi6* KO#1 and *Piwi6* KO#2 U4.4 cells were seeded in 24-well plates at 50-60% confluency and incubated for 3 days to reach a 90% confluency. A stock solution of Actinomycin-D (Act-D, Sigma; 2 mg/mL in water) was diluted in culture medium and added to the wells to a final concentration of 10 μg/mL. The treatment was performed in the dark. An equivalent amount of water was added to the wells as mock treatment. Samples were collected at 30, 45, 60, 90 and 150 min after Act-D incubation, and the control sample was collected at 30 min after the mock treatment with water. At the indicated time points, cells were harvested by trituration and pelleted by centrifugation at 285 x *g* for 5 min at 4 °C, after which the cell pellets were lyzed in 1 mL of RNA-solv.

### Rescue assays

*Piwi6* KO#1 cells were seeded in 24-well plates cells at 70–80% confluency and incubated for 16–24 h, after which they were transfected with plasmids encoding GFP-tagged wild-type *Piwi6*, the IDR deletion construct (ΔIDR), and the *Piwi6* mutants AS* and YK*. pUb-GW-GFP plasmid [32] expressing GFP was used as a negative control. For each condition, 1 μg of plasmid was transfected using 2 μL X-tremeGENE HP DNA Transfection Reagent (Roche) according to the manufacturer’s instructions. Samples were harvested 48 hours post-transfection for RNA isolation, reverse transcribed, and used in subsequent qPCR reactions to measure *AalERV1* expression.

### Protein extraction

*Piwi6* KO#1 U4.4 cells were seeded in 6-well plates and incubated for 16 hours, after which 2.5 μg of plasmids encoding GFP-tagged wild-type, ΔIDR, or mutant *Piwi6* were transfected using 5 μL X-tremeGENE HP DNA Transfection Reagent (Roche) according to the manufacturer’s instructions. At 48 hours post transfection, cells were detached by flushing the wells with 2 mL ice cold PBS. The cell suspension was then centrifuged at 1100 x *g* for 5 min at 4 °C, and the pellet was lysed in RIPA lysis buffer (10 mM Tris-HCl pH 7.4, 150 mM NaCl, 0.5 mM EDTA, 0.1 % SDS, 1% Triton X-100, 10% glycerol, 1% sodium deoxycholate [DOC] and 1x protease inhibitors [Roche]). The lysates were incubated for 1 hour at 4 °C under rotation to allow solubilization. The lysates were centrifuged at 18,000 x *g* for 30 min at 4 °C to pellet the cell debris, and the soluble fraction was taken as protein extract. Protein extracts from WT, *Piwi6* KO#1, and *Piwi6* KO#2 U4.4 cells were obtained the same way, without transfections, to analyze endogenous Piwi6 expression.

### Western Blotting

Protein samples were denatured in 2x SDS sample buffer (Bio-rad) at 95 °C for 10 min and used in SDS-PAGE and western blotting. The membranes were blocked with 5% milk powder in PBS for 1 hour at room temperature. Primary antibodies, rat-anti-Tubulin (MCA78G, Sanbio, 1:1000 dilution), rat-anti-GFP (1:1000, Chromotek, 3H9, RRID:AB_10773374), or rabbit-anti-Piwi6 (described previously [28], used at 1:500 dilution) were diluted in 2.5% milk powder in PBS and incubated overnight at 4 °C. The next day, the membranes were washed three times with PBS-T (0.1% (v/v) Tween 20 in PBS), followed by three brief PBS washes. Secondary antibodies conjugated to a fluorescence dye (IRDye 800CW goat-anti-rabbit, IRDye 680LT goat-anti-rat, Li-Cor, both at 1:10,000 dilution) were diluted in 2.5% milk in PBS-T containing 0.01% SDS and incubated for 45 min at room temperature in the dark. The membranes were washed three times with PBS-T and three times briefly with PBS and imaged on an Odyssey CLx near-infrared fluorescence imaging system (LI-COR).

### Immunofluorescence microscopy

Coverslips were cleaned overnight with 1 M HCl and washed at least three times with water, after which they were air-dried and covered completely with 50 µl of 0.5 mg/ml concanavalin A (Sigma). After an overnight incubation at room temperature, the coverslips were UV sterilized in 24-well plates for 90 sec and subsequently rinsed with PBS before cell seeding. *Piwi6* KO#1 U4.4 cells were seeded at 70-80% confluency on concanavalin A coated coverslips in 24-well plates. After 16–24 h incubation, 1 μg of plasmids encoding GFP-tagged proteins were transfected using 2 μL X-tremeGENE HP DNA Transfection Reagent (Roche) per well, following the manufacturer’s instructions. At 48 h after transfection, cells were washed with PBS and fixed using 4% paraformaldehyde (PFA). Subsequently, cells were permeabilized with 0.25% Triton X-100 in PBS, blocked with 10% Normal Goat Serum (NGS), 0.3 M Glycine, 0.1% Tween-20 in PBS, and incubated with Alexa Fluor 568 phalloidin (Invitrogen) to stain the actin cytoskeleton. After three brief PBS washes, coverslips were mounted onto slides with ProLong Gold Antifade Reagent containing DAPI (Invitrogen). All staining steps were conducted at room temperature in the dark. Slides were stored at 4 °C until imaging. Confocal images were acquired using the Zeiss LSM900 microscope, and ImageJ/Fiji was used for image processing and analysis.

Fluorescent signal intensity was quantified in the cytoplasm and nucleus of approximately 10 cells by manually tracing the phalloidin staining and DAPI staining in ImageJ/Fiji, respectively. Corrected total cell fluorescence (CTCF) (described at https://theolb.readthedocs.io/en/latest/imaging/measuring-cell-fluorescence-using-imagej.html) was calculated using the measure of *area* and the *integrated intensity*, and the following formula:

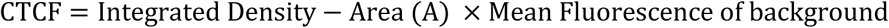

The ratio between nuclear (N) and cytoplasmic (C) fraction using the following formula:

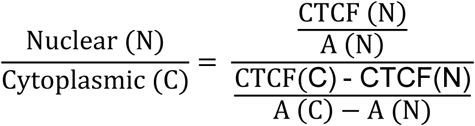

### Double stranded RNA (dsRNA) production

dsRNA was produced by T7 RNA polymerase-mediated *in vitro* transcription, using T7 promoter-flanked PCR products as templates as described [28]. PCR was performed using GO-Taq Flexi DNA Polymerase (Promega) using primers listed in Supplementary Table S4. The *in vitro* transcription reaction was incubated for 4 h at 37 °C, followed by a 10 min incubation at 95 °C and a gradual cool-down to room temperature. dsRNA was purified using the GenElute Mammalian Total RNA Miniprep Kit (Sigma). For gene knockdown in adult mosquitoes, the purified dsRNA was subsequently concentrated by ethanol precipitation.

### RNAi knockdown in cells

Wild-type U4.4 cells were seeded in 24-well plates at 50–60% confluency, and incubated for 16–24 h, after which 150 ng dsRNA was transfected using 0.3 μL of X-tremeGENE HP DNA Transfection Reagent (Roche) per well, following the manufacturer’s instructions. The same procedure was repeated at 48 hours after the first transfection to enhance knockdown efficiency. Samples were harvested in RNA-Solv Reagent (Omega Bio-tek) at 24 hours after the second knockdown. dsRNA against the *luciferase* gene was used as a non-targeting control.

### Phylogenetic analysis of PIWI proteins

Amino acid sequences of PIWI proteins were retrieved from VectorBase, FlyBase and Ensembl Metazoa (accession numbers are provided in Supplementary Table S10). Multiple sequence alignments were made using Clustal Omega and visualized with ESPript3 [34]. Sequence identity and similarity scores were calculated using Sequence Manipulation Suite [35] with multiple sequence alignment FASTA files as input. Phylogenetic trees were visualized using iTol.

### IDR and NLS prediction

The Piwi6 amino acid sequence was retrieved from VectorBase (*AALFPA_065618*) [36] and used as input for NLS and IDR prediction tools. To predict IDR, IUPred3 [37] was used with long disorder as an analysis type with medium smoothing. NLS1’ and NLS1 were predicted using NLStradamus with the threshold set to 0.6 and 0.3, respectively. A bipartite NLS (NLS2) was predicted using cNLS mapper using the default settings [38].

### Immunoprecipitation

WT U4.4 cells (clone P2B6) were seeded in 6-well plates, and four wells per condition were transfected with plasmids encoding GFP-Piwi6 (WT and YK*) or GFP the following day using X-tremeGENE HP DNA transfection reagent. At 48 h post transfection, the cells were washed twice in PBS and lysed in 250 µL ice cold RIPA buffer (10 mM Tris-HCl pH 7.5, 150 mM NaCl, 0.5 mM EDTA, 0.1% SDS, 1% Triton X-100, 1% deoxycholate, 2.5 mM MgCl_2_, 1x cOmplete protease inhibitor, 1 mM PMSF, 100 U/ml DNase I). Lysates were incubated on ice for 30 min and resuspended by pipetting every 5 min. Lysates were cleared by centrifugation at 17,000 x *g* for 10 min at 4 °C and the cleared lysate was diluted in 300 µL dilution buffer (10 mM Tris-HCl pH 7.5, 150 mM NaCl, 0.5 mM EDTA, 1x cOmplete protease inhibitor, 1 mM PMSF). For immunoprecipitation, diluted lysates were first incubated with equilibrated Binding Control magnetic agarose beads (ChromoTek) for 30 min at 4 °C with end-over-end rotation. After magnetic separation to remove the control beads, the lysates were incubated with equilibrated GFP-Trap magnetic agarose beads (ChromoTek) for 90 min at 4 °C with end-over-end rotation. The beads were washed three times in 500 µL wash buffer (10 mM Tris-HCl pH 7.5, 150 mM NaCl, 0.5 mM EDTA, 0.05% NP-40, 1x cOmplete protease inhibitor, 1 mM PMSF). RNA was extracted from the beads using 20 µg proteinase K (Ambion) in 100 μL proteinase K buffer (50 mM Tris-HCl pH 7.5, 100 mM NaCl, 1 mM EDTA, 0.5% SDS) for 90 min at 55 °C with shaking at 800 rpm, followed by RNA isolation using TRI Reagent (Sigma). Purified RNA was used as input for NEBNext Small RNA Library Prep kit (NEB) following the manufacturer’s recommendations.

### Small RNA sequencing and analysis

WT, control and *Piwi6* KO U4.4 cells were seeded in 24-well plates. Cells were either mock-infected or infected with recombinant SINV-GFP at an MOI of 1 for 48 h. Total RNA was extracted and used as input for the preparation of small RNA libraries using the NEBNext Small RNA Library Prep Set for Illumina (New England Biolabs), according to the manufacturer’s instructions as described in [28]. Libraries were pooled and sequenced on an Illumina HiSeq 4000 instrument by GenomEast Platform (Strasbourg, France). Sequence data have been deposited in the NCBI sequence read archive, BioProject accession number PRJNA989754 (for WT and control cells) and PRJNA1173503 (for *Piwi6* KO cells).

Reads were trimmed to remove the 3’ sequencing adapters using cutadapt [39] and the trimmed reads were mapped non-uniquely to the *Ae. albopictus* genome Aalbo_primary1 (RefSeq GCF_006496715.1) [40] and to the *Ae. albopictus* repeat annotation [41] with zero mismatches, and to the SINV-GFP genome allowing one mismatch using Bowtie (version 625 0.12.7) [42]. Sequencing reads mapping to annotated miRNA genes [40] were counted using bedtools [43] and used for read normalization. Results were visualized using ggplot2 (version 3.5.1) in R (version 4.4.1) [44]. Sequence logos of piRNAs mapping to transposons and to SINV-GFP genome were created using Weblogo3 [45] on 25–30 nt reads trimmed to the first 20 nt with FASTX-Toolkit (available at http://hannonlab.cshl.edu/fastx_toolkit/download.html). BedGraph files were generated from 25–30 nt RNA reads mapping to the genome with miRNAs as normalization factor using bedtools genomecov (version 2.27.1) [43], and visualized in Integrative Genomics Viewer (IGV) [46].

The GFP-Piwi6 immunoprecipitated small RNA libraries were paired-end sequenced on an Illumina NextSeq 500. The R1 reads were used for analysis. Adapter sequences were clipped, and reads were mapped to the *Ae. albopictus* reference genome Aalbo_primary1 (RefSeq GCF_006496715.1) using Bowtie, allowing no mismatches. Mapping reads were converted to BED and intersected to *Ae. albopictus* miRNA genome coordinates [34] using bedtools intersect. Clipped reads were also mapped to the *Ae. albopictus* repeat annotation using Bowtie, allowing 1 mismatch, and normalized to total miRNA counts. Sequence data have been deposited in NCBI SRA under BioProject accession number PRJNA1470531.

### mRNA sequencing library preparation and analysis

Total RNA was isolated from WT, control and *Piwi6* KO U4.4 cells in three independent experiments using RNAsolv reagent following the standard protocol. Polyadenylated mRNAs were extracted, and libraries were prepared using the TruSeq stranded mRNA Library Prep kit (Illumina) following the manufacturer’s instructions. Libraries were paired-end sequenced on an Illumina HiSeq 4000 instrument (2 x 100 bp). Sequence data have been deposited in the NCBI sequence read archive, BioProject accession number PRJNA989754 (for WT and control cells) and PRJNA1173503 (for *Piwi6* KO cells).

To quantify TE expression, RNA-seq reads were mapped to the *Ae. albopictus* repeat annotation [41] using standard settings and specifying ‘ISR’, and quantified using Salmon (version 1.5.2) [47]. Differential expression analyses and statistical testing were performed using DESeq2 (version 1.34.0) [48]. Statistical significance of differential expression between *Piwi6* KO and control cells was tested using an FDR of 0.05 without specifying a minimal log_2_ fold change. Results were visualized using ggplot2 (version 3.5.1) in R (version 4.4.1) [44] using a pseudo count of 0.5 for TEs to plot values of zero. As an additional control, a similar analysis was performed to analyze differential expression between *Piwi6* KO and WT cells with the same settings and cutoffs. To visualize RNA-seq reads in Integrative Genomics Viewer (IGV) [46], bedGraph files were generated from reads mapping to the genome using genomeCoverageBed [43] using the DEseq2 scaling factor as normalization factor.

### *AalERV1* sequence analysis

The FASTA sequence of the originally identified *AalERV1* locus (NW_021837045.1_146399653-146407556) was retrieved from the *Ae. albopictus* genome Aalbo_primary1 (RefSeq GCF_006496715.1) [40] and used as a query in a BLASTn search against the same genome. The results were filtered to retain only the regions that aligned to the query with a minimum length of 7000 nucleotides. This search yielded 28 loci with the coordinates shown in Supplementary Table S8. An *AalERV1* consensus sequence was generated using EMBOSS Cons [49] using the multiple sequence alignment of FASTA sequences from these 28 *AalERV1* loci as input.

To find *AalERV1* in the *Ae. aegypti* genome, the same query sequence was used in a BLASTn search against the TEfam transposon database (https:// tefam.biochem.vt.edu/tefam/get_fasta.php, downloaded in April 2017). The results were filtered according to the alignment length, identifying TF000419|Ty3_gypsy_Ele122 and TF000477|Ty3_gypsy_Ele134 as the top hits (alignment length of 2329 and 1271 bp, respectively).

FASTA sequences of regions corresponding to regions in Supplementary Table S8 were retrieved from the genome and used in ORFfinder [50] (https://www.ncbi.nlm.nih.gov/orffinder/) using default settings with “ATG” and alternative initiation codons option as ORF start codon, and with minimal ORF length (nt) set to 300. The translated amino acid sequences were used as query in BLASTp searches against the UniProtKB/Swiss-Prot database to find similar sequences.

### Precision Run-on Sequencing library preparation

Unless otherwise noted, all Precision Run-On sequencing (PRO-seq) steps were carried out as described in [51] with the following adaptations: for chromatin isolation, approximately 35 million WT (clone P2B6), *Piwi6* KO#1, and *Piwi6* KO#2 U4.4 cells were washed with ice-cold PBS, harvested with a cell scraper, and pelleted by centrifugation at 300 x *g* for 5 min at 4 °C. After a second wash with ice-cold PBS, resuspension in ice-cold NUN buffer, and pelleting the cells, the samples were washed with Tris-HCl pH 7.5 until a clean white chromatin pellet was obtained (two washes for WT cells and three washes for *Piwi6* KO cells). The chromatin was resuspended in chromatin storage buffer, incubated on ice for 5 min, flash frozen in liquid nitrogen, and stored at -80 °C.

Prior to the run-on reaction, chromatin samples were fragmented by sonication (Diagenode, Bioruptor Pico) using the following sequential cycles: 5 cycles of 30 sec on/30 sec off intervals at high intensity, 5 cycles of ultra-high intensity, and a final 5 cycles at high intensity. The 2X Run-On Master Mix (ROMM: 10 mM Tris-HCl pH 8.0, 5 mM MgCl_2_, 1 mM DTT (Merck Millipore), 300 mM KCl, 40 µM Biotin-11-UTP (Revvity), 40 µM ATP (NEB), 40 µM CTP (NEB), 40 µM GTP (NEB), 0.8 U/µL RNase inhibitor (Thermo Scientific), 1% Sarkosyl (Sigma-Aldrich]) was prepared fresh and pre-warmed to 30 °C. Chromatin from human Huh7 cells was added as a spike-in control. The run-on reaction was initiated by adding an equal volume of ROMM to the chromatin, mixing thoroughly by trituration, and incubated at 30 °C for 5 minutes.

Nascent transcripts were isolated, and PRO-seq libraries were prepared with barcoded samples processed individually. 5’ de-capping and 5’ hydroxy-repair were performed at 30 °C for 45 minutes, followed by 5’ adapter ligation (Supplementary Table S6), as described [51]. Reverse-transcription was performed with SuperScript IV Reverse Transcriptase (Thermo Fisher), and the resulting cDNA was amplified with Q5 Hot Start High-Fidelity DNA polymerase (NEB) with primers RP1 and RPI-n (Supplementary Table S6) under the following cycling conditions: 95° C for 2 min; 5 cycles of 95 °C for 30 sec, 56 °C for 30 sec and 72 °C for 30 sec; 11 cycles of 95 °C for 30 sec, 65 °C for 30 sec and 72 °C for 30 sec; final extension at 72 °C for 10 min. Libraries were cleaned up and size-selected using 1.4X volume AMPure XP beads (Beckman Coulter Diagnostics). Paired-end sequencing was performed on a NextSeq 500 System (Illumina) according to the standard Illumina protocol. Sequence data have been deposited in the NCBI sequence read archive, BioProject accession number PRJNA1470531.

### Precision Run-On sequencing analysis

Quality checks for raw sequencing reads were performed with FastQC (version 0.11.7) (available at https://www.bioinformatics.babraham.ac.uk/projects/fastqc/). Adapter and UMI sequences were trimmed using fastp (version 0.23.4) [52], after which the 3’ barcode was removed from the start of read 1 with fastx_trimmer (part of FASTX Toolkit version 0.0.13) and, in case of readthrough, from the end of read 2 with cutadapt (version 5.2) [39]. The reverse complement of read1 was then obtained with fastx_reverse_complement. The reads were mapped to the *Ae. albopictus* genome Aalbo_primary1 (RefSeq GCF_006496715.1) [40] and the *Ae. albopictus* repeat annotation [41] using Bowtie2 (version 2.5.4) [53] (options: --end-to-end --very-sensitive --ff --dovetail). For genome mapping, reads were sorted and indexed using SAMtools (version 1.20) [54], PCR duplicates were removed using UMI-tools (version 1.1.6) [55], and the remaining reads were filtered to retain only pairs mapping to the same chromosome. For visualization purposes, replicates were merged and RPM-normalized. Bedgraph files were created using bedtools genomecov (version 2.27.1) [43], converted to TDF files using igvtools toTDF (version 2.17.3), and visualized in IGV [46].

For mapping to repeat sequences, the *Ae. albopictus* repeat annotation was updated by replacing the *LTR/Gypsy family 3394* entry with the *AalERV1* consensus sequence generated by EMBOSS Cons [49]. PRO-seq reads were mapped to this custom repeat annotation using Bowtie2 (version 2.5.4) with the same settings as used for genome mapping. Reads were sorted and indexed using SAMtools (version 1.20), deduplicated with UMI-tools (version 1.1.6), and the number of reads mapped to each repeat annotation were extracted using idxstats from SAMtools [54]. Differential expression analysis was performed using DESeq2 (version 1.50.2) [48], with statistical significance assessed using an FDR threshold of 0.05 without specifying a log_2_ fold change cutoff. Results were visualized using ggplot2 (version 4.0.1) [44], applying a pseudo count of 0.5 to plot values of zero.

### CUT&Tag library preparation

The CUT&Tag protocol was adapted from [56], with the following modifications: WT (clone P2B6), *Piwi6* KO#1, and *Piwi6* KO#2 U4.4 cells were counted, washed once with wash buffer (20 mM HEPES pH 7.5, 150 mM NaCl, 0.5 mM spermidine (MP Biomedicals), 1x Protease Inhibitor Cocktail EDTA-free (Roche)), and transferred into digitonin wash buffer (20 mM HEPES pH 7.5, 150 mM NaCl, 0.5 mM spermidine, 1x Protease Inhibitor Cocktail EDTA-free, 0.05% digitonin (MedChemExpress)). For bead binding, 10 µL of BioMag Plus Concanavalin A beads (Bangs Laboratories, Inc.) in binding buffer (20 mM HEPES pH 7.5, 10 mM KCl, 1 mM CaCl_2_, 1 mM MnCl_2_) was added to 100,000 cells per condition and incubated for 15 min at room temperature. For primary antibody binding, cells were incubated overnight with 1 µg of antibody targeting either H3K9me3 (Diagenode, C15410193, RRID:AB_3719921) or RNA pol II CTD S5P (Abcam, ab5131, RRID:AB_449369) in antibody buffer (20 mM HEPES pH 7.5, 150 mM NaCl, 0.5 mM spermidine, 0.05% digitonin, 2 mM EDTA, 1x Protease Inhibitor Cocktail EDTA-free) on a nutator mixer at 4 °C.

The following day, samples were washed twice with digitonin wash buffer before secondary antibody binding. Cells were resuspended in antibody buffer supplemented with 1 µg Donkey Anti-Rabbit IgG H&L (Abcam, ab6701, RRID:AB_956011) and incubated at 4 °C for 1 h. After two washes with digitonin wash buffer, pAG-Tn5 (EpiCypher) was bound by adding 150 µL digitonin high salt wash buffer (20 mM HEPES pH 7.5, 300 mM NaCl, 0.5 mM spermidine, 0.05% digitonin, 1x Roche Protease Inhibitor Cocktail EDTA-free) supplemented with 2.5 µL pAG-Tn5 and incubating for 1 hour at room temperature on a nutating mixer.

For chromatin digestion and clean-up, cells were washed twice with digitonin high salt wash buffer and resuspended in tagmentation buffer (20 mM HEPES pH 7.5, 300 mM NaCl, 0.5 mM spermidine, 10 mM MgCl_2_, 0.05% digitonin, 1x Protease Inhibitor Cocktail EDTA-free), then incubated shaking at 700 rpm in a ThermoMixer C (Eppendorf) for 1 h at 37 °C. The reaction was stopped by addition of 22.5 mM EDTA, 0.5% SDS, and 0.4 µg/µL Proteinase K (Invitrogen) (final concentrations), followed by incubation at 700 rpm for 30 min at 55 °C, and a further 20 min at 70 °C. DNA was purified using the ChIP DNA Clean & Concentrator kit (Zymo Research) following the manufacturer’s instructions and size-selected using 2.5x volume AMPure XP beads (Beckman Coulter Diagnostics).

Libraries were prepared by supplementing 2x NEBNext Ultra II Q5 Master Mix and indexed with Illumina i5 and i7 primers (Supplementary Table S7) using the following cycling conditions: 72 °C for 5 min; 98 °C for 30 sec; 11 cycles (H3K9me3) or 14 cycles (RNA Pol II) of 98 °C for 10 sec and 63 °C for 30 sec; final extension at 72 °C for 1 min. Libraries were cleaned-up by two rounds of purification using 0.9x volume AMPure XP beads, followed by a two-sided size-selection of 0.55x and 1.5x volume AMPure XP beads. Samples were paired-end sequenced on a NextSeq 500 System (Illumina) according to the standard Illumina protocol. Sequence data have been deposited in the NCBI sequence read archive, BioProject accession number PRJNA1470531.

### CUT&Tag analysis

The CUT&Tag analysis was adapted from [57]. Quality checks for raw sequencing reads were performed with FastQC (version 0.11.7) (available at https://www.bioinformatics.babraham.ac.uk/projects/fastqc/). Reads were mapped to the *Ae. albopictus* genome Aalbo_primary1 (RefSeq GCF_006496715.1) [40] or *Ae. albopictus* repeat annotation [41] using Bowtie2 (version 2.5.4) [53] (options: --end-to-end --very-sensitive --no-mixed --no-discordant –phred33 -I 10 -X 700). For genome mapping, reads were sorted by coordinate, indexed, and deduplicated using Picard (version 3.3.0) (available at https://broadinstitute.github.io/picard/). Mapped read pairs were then filtered to remove unmapped reads using SAMtools (version 1.20) [54], followed by a second filtering to retain only read pairs mapping to the same chromosome with fragments length below 1000 bp. For visualization purposes, replicates were merged and RPM-normalized bedgraph files were created using bedtools genomecov (version 2.27.1) [43], converted to TDF files using igvtools toTDF (version 2.17.3) and visualized in IGV [46] .

For repeat annotation mapping, CUT&Tag reads were mapped to the custom repeat annotation described for PRO-seq using Bowtie2 (version 2.5.4) with the same settings as used for the genome mapping. Reads were sorted and indexed, and the number of reads mapping to each repeat were extracted using idxstats from SAMtools (version 1.20). Differential expression analyses and visualization were performed as described for the PRO-seq data. For the violin plots, data normality was tested using the Shapiro-Wilk test with shapiro.test (in R package stats, version 4.5.2), followed by either a repeated-measures ANOVA using anova_test with pairwise paired t-tests using pairwise_t_test or a Friedman test using friedman_test with pairwise Wilcoxon signed-rank test using wilcox_test (in R package rstatix, version 0.7.3) (available at https://doi.org/10.32614/CRAN.package.rstatix), depending on normality.

### Mosquito Rearing

*Ae. albopictus* mosquitoes (AGROPOLIS strain) were reared under standard insectary conditions. The mosquitoes were housed in a secure rearing facility maintained at 27 °C (+/-1°C), at 70% (+/- 20%) humidity, and 16:8 h light/dark cycle. The mosquitoes were provided with 6% sucrose and were fed with bovine and human blood using the Hemotek PS5 feeder (Discovery Workshops, UK) with 3 ml reservoirs (FU1-3) with a parafilm membrane.

### *Ae. albopictus* tissue dissections

Three- to five-day-old, non-blood fed female mosquitoes were used to measure PIWI gene expression in tissues. Approximately 20 mosquitoes were dissected in PBS, and individual tissues were pooled in 1.5 ml screw cap tubes containing 1.0 mm zirconia beads (Biospec Products) and 50 μL PBS. The dissected tissues were kept on ice until 1 mL of RNA-solv reagent (Omega Bio-Tek) was added to each tube. Tissues were homogenized in RNA-solv reagent using a Precellys 24 homogenizer (Bertin Technologies). Subsequently, RNA was isolated, reverse transcribed and used as template in end-point PCR. Primers used in the PCR reactions are shown in Supplementary Table S5.

### *Piwi6* knockdown in adult *Ae. albopictus* mosquitoes

*Piwi6* gene knockdowns were performed through intrathoracic injections using three-to five-day-old, non-blood fed female mosquitoes. The mosquitoes were immobilized on ice, and either 240 ng of dsRNA against *Piwi6* or dsRNA against the *Luciferase* gene was injected per mosquito using a Nanoject III microinjector (Drummond). Approximately 50 mosquitoes per condition were injected. Two days after the injections, mosquitoes were starved by removing the sugar water, and the following day, mosquitoes were blood fed through arm feeding. Only fully engorged mosquitoes were retained for subsequent steps. Four days after the blood meal, mosquito ovaries were dissected and the remaining parts were retained as carcasses. Individual ovaries and carcasses were placed in 1.5 ml screw cap tubes containing 1.0 mm zirconia beads (Biospec Products) and 50 μL PBS and kept cold until RNA extraction. 1 mL of RNA-solv reagent (Omega Bio-Tek) was added to the tubes and the tissues were homogenized using a Precellys 24 homogenizer (Bertin Technologies).

### Statistical Analysis

Statistical analyses were performed using GraphPad Prism9 software and R statistics version 4.5.2. Differences were tested for statistical significance using unpaired two-tailed *t*-tests, one-way ANOVA with Holm-Sidak’s multiple comparisons post hoc test, repeated-measures ANOVA with pairwise paired *t*-tests or Friedman tests with pairwise Wilcoxon signed-rank tests, as specified. Data normality of CUT&Tag data was tested using the Shapiro-Wilk test.

## Results

### *Piwi6* is broadly expressed in gonads and somatic tissues of *Aedes albopictus*

PIWI proteins in *Ae. albopictus* are highly similar to those of *Ae. aegypti* and share the overall domain structure with their orthologs in the fruit fly *Drosophila melanogaster* (Piwi) and the silk moth *Bombyx mori* (Siwi). Specifically, *Ae. albopictus* Piwi6 is > 90% identical to *Ae. aegypti* Piwi6 at the amino acid level, while it has around 55% similarity and 40% identity to Piwi and Siwi (Figure 1A-B). Moreover, experimentally validated residues with crucial functions, such as piRNA binding sites and catalytically active sites are conserved in *Aedes* Piwi6 (Figure 1C, Figure S1). We evaluated the expression pattern of PIWI genes in dissected tissues from non-blood fed, female *Ae. albopictus* mosquitoes by RT-PCR. *Piwi4*, *Piwi6* and *Ago3* were broadly expressed in legs, heads, midguts, salivary glands, Malpighian tubules, carcasses and ovaries, while *Piwi5* was highly expressed in carcasses and ovaries, lowly expressed in legs, heads, salivary glands and Malpighian tubules, and undetectable in midguts (Figure 1D). In contrast, *Piwi2* and *Piwi3* were expressed in the ovaries and carcasses, and *Piwi7* was undetectable in any adult tissue. These results confirm and extend previous observations in *Aedes aegypti* that the expression of members of the expanded PIWI family is not restricted to the gonads [14,15].

### Piwi6 is not required for the biogenesis of virus and host derived piRNAs

We used CRISPR/Cas9 engineering to generate *Piwi6* knockouts in the *Ae. albopictus* U4.4 cell line. Two independent clones contained frameshift mutations leading to premature stop codons in all *Piwi6* alleles (Supplementary Figure S2A), which was confirmed by RT-qPCR analysis showing a strong reduction in *Piwi6* mRNA expression (Supplementary Figure S2B), likely as a result of nonsense-mediated RNA decay. Indeed, Piwi6 protein was undetectable in western blot analysis, confirming the knockout phenotype of these cells (Figure 2A). Overall, microRNA and endogenous piRNA counts, total and nascent mRNA, and TE expression correlated well between both *Piwi6* KO clones (Supplementary Figure S3), and we therefore used the average read counts from both clones throughout this study. In parallel to the *Piwi6* KO cells, we used *Cas9* transfection in the absence of gene-specific guide RNAs to generate control cells (Ctrl), which were used along with the parental wild-type (WT) U4.4 cells in some experiments.

Previous studies have found that viral RNA is processed into viral (v)piRNAs in *Aedes* cell lines and adult mosquitoes upon infection with arthropod-borne viruses [27,58–60]. To assess whether Piwi6 plays a role in vpiRNA production, we infected *Piwi6* KO and WT cells with recombinant Sindbis virus expressing GFP (SINV-GFP) and analyzed small RNA expression by RNA sequencing. While the size distribution of small RNAs mapping to the SINV-GFP genome did not change in *Piwi6* KOs compared to WT cells, the number of SINV-derived piRNA reads was reduced in *Piwi6* KO cells, which could not be explained by differences in viral RNA replication (Supplementary Figure S2C-S2D). This reduction in vpiRNA levels was relatively minor compared to the effect observed for *Piwi5* KO cells, in which vpiRNA production was completely abolished [62]. Moreover, sequence logos of the remaining vpiRNAs were unaffected by Piwi6 loss, with strong enrichments for a uridine at position 1 of antisense piRNAs and an adenosine at position 10 of sense piRNAs, indicative of a functional ping-pong amplification cycle (Supplementary Figure S2E). These results argue against an essential role for Piwi6 in vpiRNA biogenesis and are in line with our earlier observations upon *Piwi6* knockdown in *Ae. aegypti* cells [25,62].

**Figure 1.**
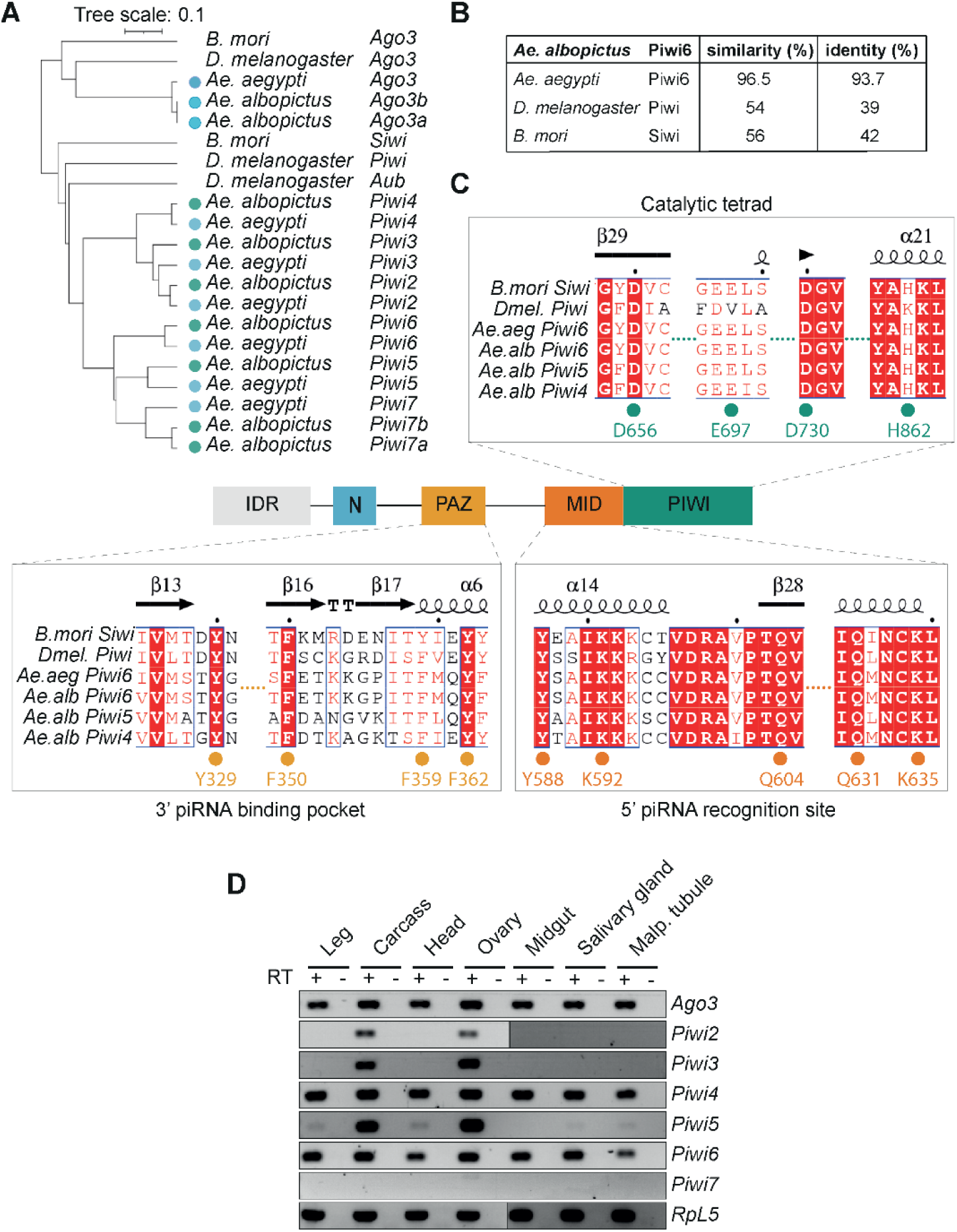
*Piwi6* is expressed in gonadal and non-gonadal tissues of *Ae. albopictus* mosquitoes. A) Unrooted neighbor joining tree based on amino acid sequences of *Ae. albopictus* Piwi6 and its orthologs in *Ae. aegypti*, *Drosophila melanogaster* and *Bombyx mori*. In the earlier version of the *Ae. albopictus* reference genome (Foshan strain, AaloF1) *Piwi1* and *Piwi3* were annotated as two separate entries, which were merged into a single gene in later reference genomes. The scale bar corresponds to 0.1 substitutions per site. B) Percent similarity and identity of the *Ae. albopictus* Piwi6 amino acid sequence to *Ae. aegypti* Piwi6, *Drosophila melanogaster* Piwi and *Bombyx mori* Siwi. C) Multiple sequence alignment of the *Aedes* Piwi4–6, *Drosophila melanogaster* Piwi and *Bombyx mori* Siwi sequences. Key residues are indicated by colored dots and the secondary structure of Siwi is indicated above the sequences [61]. D) PIWI gene expression in the indicated tissues from non-blood fed female *Ae. albopictus* mosquitoes, analyzed by RT-PCR. The housekeeping gene *60S ribosomal protein L5 (RPL5)* was used as a positive control. cDNA synthesis was performed in the presence (+) or absence (-) of reverse transcriptase (RT) to control for contamination of RNA preparations with chromosomal DNA. Malp. tubule, malpighian tubule; carcass, the remaining parts of the animal after tissue dissection. Uncropped gel images are shown in Supplementary Figure S10. See also Figure S1 and S10.

We next mapped the small RNA sequences to the *Ae. albopictus* genome to study how Piwi6 loss affects the expression of endogenous small RNAs. While the levels of 21–24 nt reads, depicting putative small interfering RNAs and microRNAs, were unaffected by Piwi6 loss, the levels of piRNA-sized reads were increased in *Piwi6* KO cells compared to WT and Ctrl cells (Figure 2B). This was unexpected as we anticipated that loss of a PIWI protein would result in reduced expression of the piRNA population bound to it [10,63]. To study which specific population of piRNAs were deregulated upon Piwi6 loss, we systematically analyzed the main classes of endogenous piRNAs derived from nrEVEs, piRNA clusters, TEs and other repeat elements. No major differences were observed in the expression of nrEVE-mapping piRNAs in *Piwi6* KO compared to WT cells, regardless of their orientation (Supplementary Figure S4A-B). We next analyzed expression from piRNA clusters, using our recent annotation containing 285 piRNA clusters in *Ae. albopictus* U4.4 cells, 37 of which ubiquitously expressed across tissues (core-clusters) [29]. Overall, no generalized loss of piRNA expression was observed. Although an increased number of piRNAs mapped to some clusters upon Piwi6 loss, this effect was relatively mild (mean 1.02- and 1.10-fold increase for core clusters and all clusters, respectively; Supplementary Figure S4C-D). Moreover, the same clusters were also affected in the Ctrl cells, suggesting a clonal effect rather than a direct effect of Piwi6 loss (Supplementary Figure S4C-D). Overall, these data suggest that Piwi6 is not generally required for the biogenesis of virus-, nrEVE-, and piRNA cluster-derived piRNAs.

### Increased expression of LTR retrotransposons upon Piwi6 loss

TEs are a major source of endogenous piRNAs, and we thus counted and categorized TE-mapping piRNAs using the RepeatMasker annotation [40] and noticed a 1.7- and 1.9-fold increase in the total number of LTR retrotransposon-derived piRNAs upon Piwi6 loss compared to WT and Ctrl cells, respectively (Figure 2C), In contrast, piRNAs mapping to other TE classes, as well as satellite repeats, simple repeats and unclassified repeat classes were unaffected (Figure 2C). Interestingly, increased expression was most prominent for piRNAs mapping in sense orientation to TEs (Figure 2D), and these sense piRNAs had a stronger 10A bias in *Piwi6* KO cells than in WT and Ctrl cells, a signature of ping-pong amplified piRNAs (Figure 2E).

**Figure 2.**
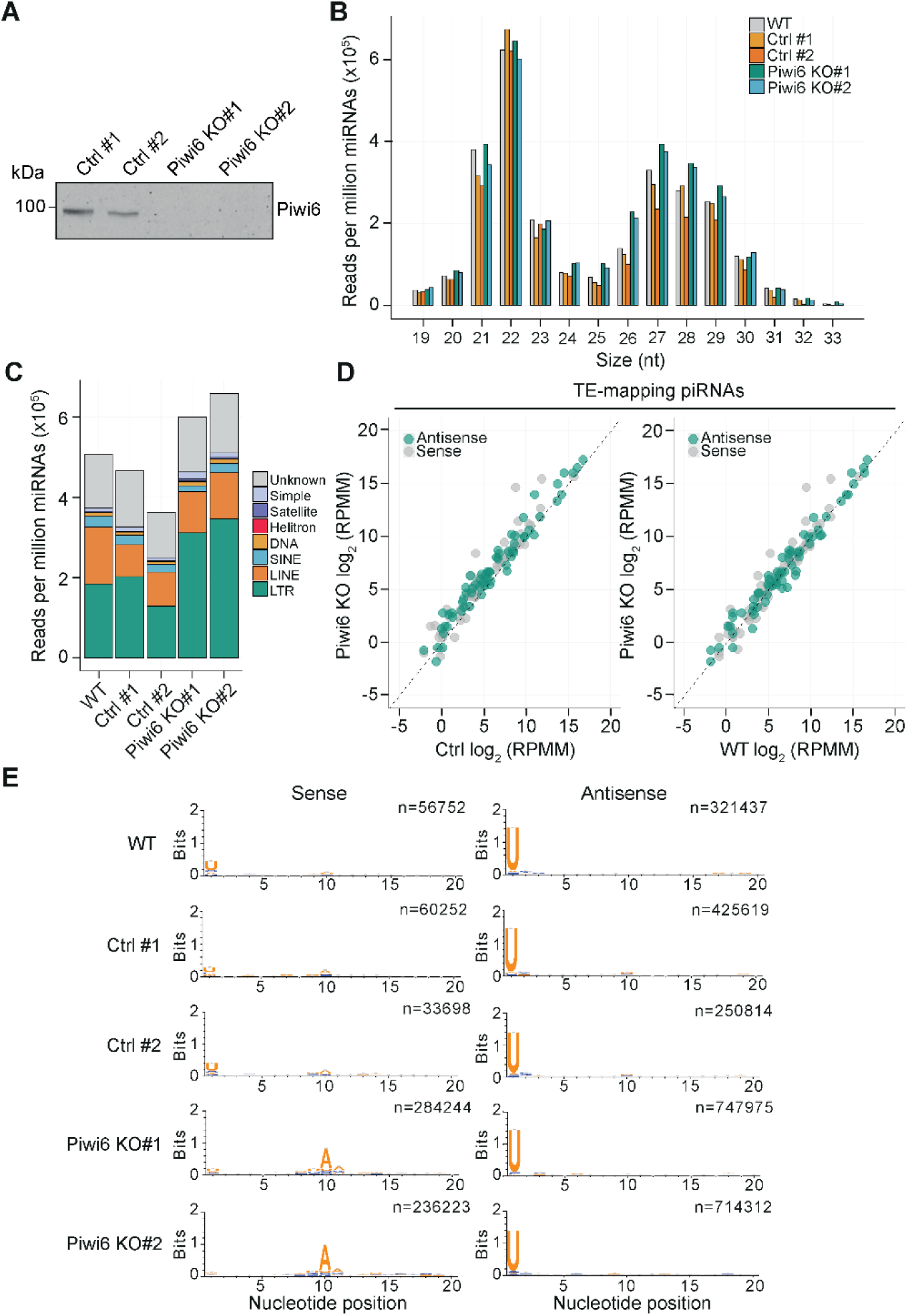
Increased mRNA and piRNA expression of LTR retrotransposons in *Ae. albopictus Piwi6* knockout cells. A) Western blot analysis of Piwi6 expression in CRISPR Control (Ctrl) and *Piwi6* knockout (KO) U4.4 cells. Uncropped gel images are shown in Supplementary Figure S10. B) Size profiles of small RNA reads mapping to the *Ae. albopictus* genome without mismatches in libraries from wild-type (WT), control (Ctrl) and *Piwi6* knockout (KO) cells. Read counts were normalized to the total number of miRNAs (in millions). C) Transposable element (TE)-derived piRNA expression in WT, Ctrl and *Piwi6* KO cells categorized according to TE order. Read counts were normalized to the total number of miRNAs (in millions). D) Scatterplots showing TE-derived piRNA expression in WT, Ctrl and *Piwi6* KO cells. piRNAs mapping to different TE orders in sense (gray) and antisense (green) orientation were counted separately, a pseudo count of one was added to plot values of zero, normalized to the number of miRNAs (in millions), and expressed as log2-transformed reads per million miRNAs (RPMM). For Ctrl and *Piwi6* KO clones the mean of two clones is shown. E) Nucleotide biases of piRNAs mapping in sense (top) or antisense (bottom) orientation to *Ae. albopictus* repeat elements in WT, Ctrl and *Piwi6* KO cells. The number of reads contributing to each sequence logo are indicated. See also Figure S2, S3, S4 and S10.

In ping-pong amplification, antisense piRNAs guide the cleavage of mRNA transcripts, which are then processed into sense piRNAs. Given the increased levels of sense piRNAs from LTR retrotransposons in *Piwi6* KO cells, we speculated that Piwi6 loss results in desilencing of these elements and increased number of transcripts that serve as substrates for piRNA production. To test this hypothesis, we performed RNA-sequencing on WT, Ctrl and *Piwi6* KO cells and mapped the reads to the *Ae. albopictus* repeat annotation [41]. *Piwi6* loss resulted in differential expression of 206 out of a total of 2533 TE sequences (adjusted *P* value ≤ 0.05; no threshold for log_2_ fold change) (Figure 3A-B). Of those, 145 elements were significantly upregulated and 61 downregulated compared to Ctrl cells, and similar results were obtained when differential expression was tested against WT instead of Ctrl cells, with 181 elements significantly upregulated and 193 downregulated compared to WT cells (Supplementary Figure S3E-F). Among the differentially expressed TEs, LTR retrotransposons were particularly affected, with 108 LTR elements significantly upregulated and 32 elements downregulated in *Piwi6* KO cells (Figure 3A, Supplementary Figure S4E). Beyond the number of differentially expressed TEs, LTR elements also showed the highest level of desilencing in *Piwi6* KO cells: 19 elements showed an increase in expression of > 16-fold (range, 16 to 826-fold; Figure 3B).

To determine whether differentially expressed TEs are potential targets of Piwi6 associated piRNAs, we performed GFP-Piwi6 immunoprecipitation followed by small RNA sequencing. piRNAs mapping to the upregulated elements, including the highly upregulated element LTR Gypsy family-3394, were enriched in GFP-Piwi6 IP compared to GFP control IP (Figure 3C, Supplementary Figure S5A). Notably, the majority of GFP-Piwi6 associated piRNAs mapped in antisense orientation to the LTR Gypsy family-3394 element, consistent with the potential role for Piwi6-bound piRNAs in transposon silencing (Supplementary Figure S5B). Together, these data indicate that Piwi6 is particularly required for silencing LTR retrotransposons, the elements with the highest abundance and rate of recent transposition in the *Ae. albopictus* genome [64].

### Piwi6 regulates a full-length endogenous retrovirus

To find out which regions in the genome were affected by Piwi6 loss, we manually inspected genomic regions and noticed a 7904 bp sequence that was strongly upregulated in *Piwi6* KO cells (Figure 3D, Supplementary Figure S5C). These upregulated transcripts seem to be processed into piRNAs, as expression of both sense and antisense piRNAs derived from this sequence was increased in *Piwi6* KO cells (Figure 4A). This locus was flanked by sequences corresponding to LTR/Gypsy family-43 in RepeatMasker at both the 5’ and 3’ ends and contained sequences corresponding to LTR/Gypsy family-210 as well as a region that was unannotated in the *Ae. albopictus* genome (Supplementary Figure S5C) [40]. Moreover, one of the TE elements that showed the highest upregulation upon *Piwi6* loss, LTR Gypsy family-3394 (22-fold; Figure 3B), aligned to this locus with high identity over two regions (96% and 99% for the first and the second region, respectively) (Supplementary Figure S5C).

The locus was predicted to encode ORFs with similarity to the *gag*, *pol* and *env* genes of infectious endogenous retroviruses, such as the gypsy element in *Drosophila melanogaster* [65,66] (Figure 3D). Specifically, the first ORF displayed similarity to the *gag* gene, the second ORF showed similarity to *pol* gene that encodes the protease, reverse transcriptase, RNase H and integrase enzymes, and the third ORF displayed similarity to gypsy *envelope* (Supplementary Figure S5C). Intriguingly, this locus was flanked by 403-bp long identical sequences, likely representing the LTRs, which are usually between 300 and 400 bp long [67,68]. The presence of intact *gag*, *pol*, and *env* genes flanked by LTR sequences, strongly suggests that this locus represents a full-length endogenous retrovirus, which we named “*AalERV1* (*<u>A</u>edes <u>al</u>bopictus* <u>E</u>ndogenous <u>R</u>etro<u>v</u>irus-<u>1</u>)”.

Using BLASTn, we identified 28 highly similar, (near) full-length *AalERV1* loci in the genome, ranging from 7007 bp to 7908 bp in size (Supplementary Table S8). All *AalERV1* loci have a minimum of 94% identity at the nucleotide level, suggesting that they are different insertion sites of the same element. Manual inspection of these *AalERV1* loci indicated that six out of these 28 insertions contain all three intact ORFs (e.g., Supplementary Figure S5C), while the others seem to be defective due to the presence of premature stop codons.

We next queried the genome of the closely related *Ae. aegypti* and found two hits, TF000477_Ty3_gypsy_Ele134 and TF000419_Ty3_gypsy_Ele122, that were 54% and 48% identical to *AalERV1*, respectively [69]. Following the convention that over 80% sequence identity is required in at least 80% of the aligned sequence for TEs to be considered members from the same family [69], we did not classify these two elements as *AalERV1*. Together, these analyses suggest that Piwi6 regulates a full-length, and therefore putatively active *Ae. albopictus* specific endogenous retrovirus.

**Figure 3.**
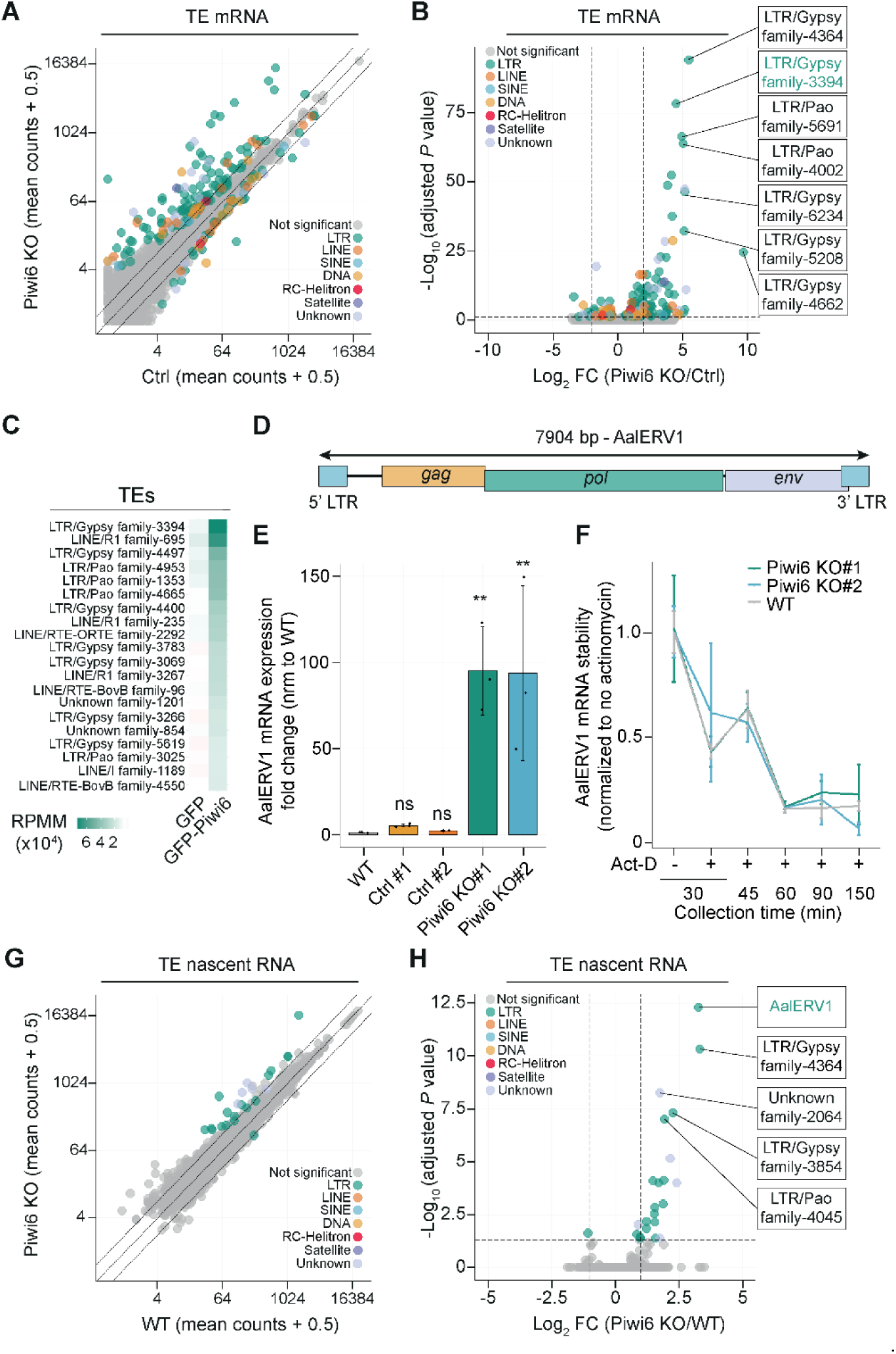
*AalERV1* is not silenced in *Ae. albopictus Piwi6* knockout cells. A) Expression of TE mRNAs in Ctrl and *Piwi6* KO cells measured by RNA-seq. Data represent mean read counts of three biological replicates. A pseudo count of 0.5 was added to all values to plot values of zero. Outer diagonal lines indicate log_2_ fold change (FC) > 1. Gray symbols, not significant; colored symbols, differentially expressed TEs categorized according to the *Ae. albopictus* repeat annotation [41]. B) Volcano plot showing the log_2_ fold changes in TE mRNA expression between *Piwi6* KO cells and Ctrl cells. Vertical dashed lines highlight log_2_FC > 2, and horizontal dashed lines highlight the threshold of statistical significance at an adjusted *P* value ≤ 0.05. Names of individual elements based on the *Ae. albopictus* repeat annotation [41] are indicated. C) Heatmap showing the top 20 TE elements enriched in GFP-tagged Piwi6 immunoprecipitation (IP) compared to GFP alone. Small RNA reads were mapped to *Ae. albopictus* repeat annotation [41] allowing one mismatch, read counts were normalized to the total number of miRNAs (in millions) (RPMM). TEs showing > 2-fold enrichment in GFP-Piwi6 over GFP IP were selected and the top 20 elements are shown, ranked by read counts in GFP-Piwi6. D) Structure of *AalERV1* comprising three open reading frames corresponding to *gag*, *pol* and *envelope*, flanked by 5’ LTR and 3’ LTR sequences. E) *AalERV1* expression in WT, Ctrl and *Piwi6* KO cells, assessed by RT-qPCR. Data represent mean ± SD of three biological replicates in an experiment representative of two independent experiments. One-way ANOVA with Holm-Sidak’s multiple comparisons post hoc test was performed to determine statistically significant differences with the WT cells, ** *P* < 0.01, ns; not significant. F) *AalERV1* mRNA stability in WT and *Piwi6* KO cells, assessed by RT-PCR at indicated timepoints after treatment with Actinomycin D (Act-D). Lines represent mean ± SD of three biological replicates in one experiment. mRNA expression was normalized to the housekeeping gene *RPL5* and expressed relative to the mock treated conditions. G) Nascent TE RNA expression in WT and *Piwi6* KO cells, as measured by PRO-seq. Data represent the mean of two biological replicates, with a pseudo count of 0.5 added to enable plotting of values of zero. Outer diagonal lines indicate log_2_FC > 1. Gray symbols, not significant; colored symbols, differentially expressed TEs categorized according to the *Ae. albopictus* repeat annotation [41]. H) Volcano plot showing the log_2_ fold changes in nascent TE RNA expression between *Piwi6* KO and WT cells. Vertical dashed lines highlight log_2_FC > 1, and the horizontal dashed line marks the threshold of statistical significance at adjusted *P* values ≤ 0.05. Individual elements are labeled according to the *Ae. albopictus* repeat annotation [41]. See also Figure S3, S4, S5.

### Piwi6 does not silence *AalERV1* at the post-transcriptional level

Using RT-qPCR, we confirmed that *AalERV1* mRNA expression was strongly increased in the two *Piwi6* KO clones compared to WT and Ctrl cells (up to 90-fold; Figure 3E). PIWI proteins may regulate gene expression at the post-transcriptional or transcriptional level [2,6,10,11]. If Piwi6 acts in post-transcriptional silencing, one would expect a change in the *AalERV1* mRNA stability upon Piwi6 loss. To study this, we blocked transcription using Actinomycin-D for 0–2.5 hours and evaluated *AalERV1* transcript turnover rates by RT-qPCR. We observed no discernible differences in decay rates in WT and *Piwi6* KO cells, indicating that Piwi6 loss does not impact *AalERV1* mRNA stability (Figure 3F). Overall, these findings strongly argue against a post transcriptional mechanism for *AalERV1* regulation by Piwi6.

### Transcriptional silencing of *AalERV1* by Piwi6

To investigate whether Piwi6 regulates *AalERV1* at the transcriptional level, we analyzed the nascent transcriptome using Precision Run-On sequencing (PRO-seq) in WT and *Piwi6* KO cells. For the data analysis, we generated an updated repeat annotation in which LTR Gypsy family-3394 element was replaced by the *AalERV1* sequence to more accurately represent the full-length element. Consistent with the RT-qPCR and RNAseq results, nascent transcript levels of *AalERV1* were strongly increased in *Piwi6* KO cells compared to WT cells (Figure 3G-H, Figure 4A and Supplementary Figure S3G). Together, these findings support a role for Piwi6 in transcriptional silencing of *AalERV1*.

In *Drosophila*, Piwi silences TEs in the ovarian somatic follicle cells at the transcriptional level by guiding the deposition of repressive chromatin marks on transposon loci, resulting in the formation of heterochromatin [9–12]. Since our PRO-seq analyses indicated that Piwi6 regulates *AalERV1* at the level of nascent transcription, we next investigated whether Piwi6 similarly affects the chromatin state at the *AalERV1* loci. To address this, we performed Cleavage Under Targets and Tagmentation (CUT&Tag) for RNA polymerase II, phosphorylated at serine 5 in the C-terminal domain (RNA pol II), a marker of active transcription, and H3 lysine 9 trimethylation (H3K9me3), a major repressive histone mark in WT and *Piwi6* KO cells. Consistent with a transcriptional silencing mechanism, *Piwi6* knockout caused a strong increase in RNA pol II occupancy at *AalERV1* (Figure 4A-B, Supplementary Figure 6C) and a depletion of H3K9me3 (Figure 4A, Figure 4C, Supplementary Figure 6B), creating a chromatin environment permissive for *AalERV1* transcription.

**Figure 4.**
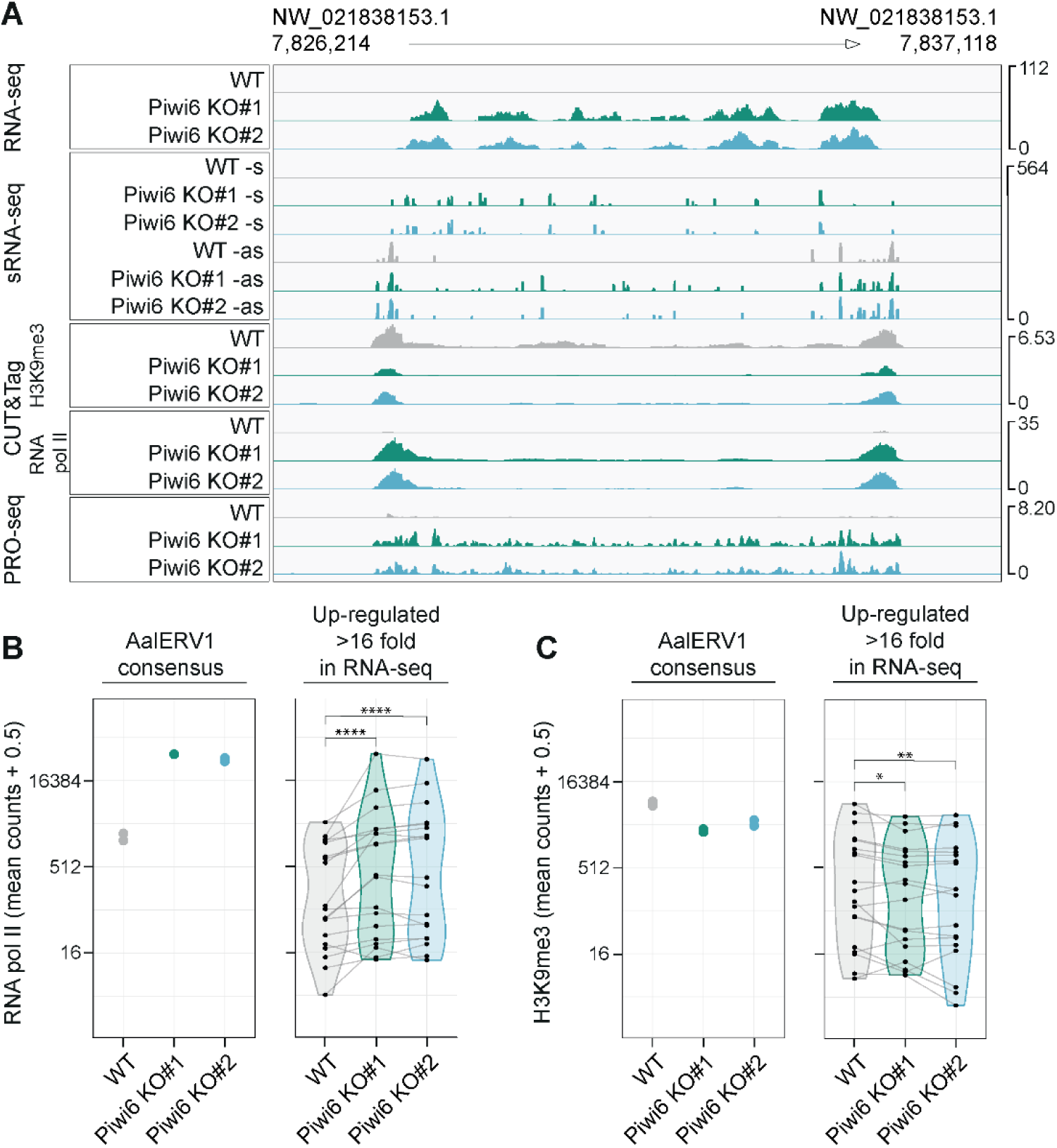
Piwi6 silences *AalERV1* at the transcriptional level. A) Genome browser tracks of a representative *AalERV1* locus (NW_021838153.1:7827714-7835618) displaying profiles of RNA seq reads, piRNA reads, CUT&Tag reads, and PRO-seq reads in WT and *Piwi6* KO cells. RNA-seq reads were normalized with DESeq2, piRNA reads were normalized to the number of miRNAs (in millions) and CUT&Tag and PRO-seq reads were RPM-normalized. B-C) Enrichment of RNA polymerase II (RNA pol II; B) and H3K9me3 (C) at TEs in WT and *Piwi6* KO cells, as measured by CUT&Tag. Reads were normalized with DESeq2. Violin plots show the distribution across TEs upregulated more than 16-fold in RNA-seq, with each data point representing the mean of two biological replicates, and lines connecting the TEs. The dot plot shows individual biological replicates for the *AalERV1* consensus sequence. A pseudo count of 0.5 was added to all plots to enable plotting of values of zero. Data in B and C were analyzed using a repeated-measures ANOVA with Bonferroni-corrected pairwise *t*-test on log2-transformed counts; * *P* < 0.05; ** *P <* 0.01; **** *P* < 0.0001 See also Figure S3, S4, S5.

To correlate RNA pol II occupancy and H3K9me3 enrichment with TE expression, we stratified TEs into four categories based on their differential RNA expression in *Piwi6* KO cells (Figure 3A and 3B): significantly upregulated more than 16-fold (range, 16 to 826-fold; n = 19), significantly upregulated (range, 1.5 to 826-fold; n = 145), significantly downregulated (range, 1.3 to 10-fold; n = 61), and not significantly deregulated (n = 2327). We then quantified RNA pol II and H3K9me3 signals over these four categories, as well as all repeats combined (Figure 4B, Figure 4C and Supplementary Figure S6A). We observed a significant increase in RNA pol II occupancy and a H3K9me3 depletion for the elements that were strongly (> 16-fold) upregulated in *Piwi6* KO cells (Figure 4B-C). In contrast, the elements with mild upregulation (1.5 to 16-fold), also showed a clear increase in RNA pol II occupancy while changes in H3K9me3 were more variable and not uniformly reduced across all elements, suggesting that increased transcriptional activity can occur independently of H3K9me3 loss for some of these elements (Supplementary Figure 6A). As expected, there were no changes in RNA pol II occupancy and H3K9me3 levels for all repeats combined, nor for elements that were not significantly regulated or downregulated in *Piwi6* KO cells (Supplementary Figure 6A). Collectively, our data support a role for Piwi6 in transcriptional silencing of *AalERV1* and a subset of highly derepressed transposable elements.

### Piwi6 localizes to the nucleus

A transcriptional mode of action requires Piwi6 to be active in the nucleus as observed in *Drosophila,* where Piwi loaded with a piRNA in the cytoplasm translocates to the nucleus to mediate transcriptional repression [70,71]. To test if Piwi6 localizes to the nucleus, we analyzed the subcellular localization of a GFP-tagged *Piwi6* transgene in *Piwi6* KO cells. We found that GFP-Piwi6 was localized throughout the cell, with a slight enrichment in the nucleus, whereas the GFP control was equally distributed between both compartments (Figure 5B, quantified in Figure 5C). The observed nuclear localization of Piwi6 is consistent with findings in *Ae. aegypti* cells [32], in which Piwi6 localizes to both the cytoplasm and nucleus and interacts with proteins that have a nuclear localization [28].

### The intrinsically disordered region of Piwi6 is required for nuclear localization

We next analyzed the determinants of Piwi6’s nuclear localization using a series of point mutants and deletion mutants (Figure 5A). We mutated aspartate 656 in the putative catalytic tetrad to alanine (DEDH > AEDH, indicated as active site: as*; Figure 1C, Figure 5A), as a single mutation in the catalytic tetrad was sufficient to abolish Siwi’s slicer activity [61]. Subcellular localization of this active site mutant resembled that of wild-type GFP-Piwi6 (Figure 5B-C), indicating that the catalytic site does not contribute to the nuclear localization of Piwi6. We further explored whether piRNA loading of impacts Piwi6 localization, as previously observed for Piwi and Aub in *Drosophila* and for MIWI2 in mice [11,18,72–75]. To investigate this, we replaced the residues in the conserved Y-K motif of the 5′ piRNA binding pocket (Figure 1C) [9,61,76] with alanine (Y588A, K592A; YK*) (Figure 5A). While we confirmed the YK* mutant to be defective in piRNA binding (Supplementary Figure 7A), the subcellular localization of this mutant was not significantly different from wild-type GFP-Piwi6 (Figure5B and C), suggesting that piRNA loading is not essential for nuclear localization, consistent with previous observations that a substantial fraction of piRNA-deficient *Drosophila* Piwi resided in the nucleus [9].

Considering that the nuclear localization signal (NLS) of *Drosophila* Piwi is located in the highly variable and intrinsically disordered N terminal region of Piwi [74,75], we speculated that the intrinsically disordered region (IDR) of Piwi6 might play a role in its nuclear localization. We deleted the entire Piwi6 IDR, as determined using the AlphaFold model prediction [77] retrieved from VectorBase [36] and IUPred3 server [37] (Figure 5A and Supplementary Figure S7B). In stark contrast to the other Piwi6 mutants tested, this IDR deletion mutant (GFP-Piwi6-ΔIDR) was clearly displaced from the nucleus to the cytoplasm (Figure 5B–C), indicating that the IDR contains the nuclear localization signal. Indeed, a fusion protein of the Piwi6 IDR with GFP (IDR-GFP) strongly accumulated in the nucleus, in comparison to GFP which was distributed throughout the cell (Figure 5D-E).

We next predicted potential NLSs in the Piwi6 IDR using NLStradamus [78] and cNLS mapper [38] (Supplementary Figure S7C) and generated deletion mutants lacking single (ΔNLS1, ΔNLS1’, ΔNLS2) or a combination of putative NLS sites (ΔNLS1+2) (Supplementary Figure S7C-D). In contrast to the NLS1 and NLS1’ deletions, the NLS2 deletion strongly reduced the nuclear localization of Piwi6 (Supplementary Figure S7D-E, Figure 5B-C). Combined deletion of NLS2 with NLS1 did not further affect Piwi6 localization, suggesting that NLS2 (residues 52-85) is a major determinant of the nuclear translocation of Piwi6. In an attempt to further map the NLS independent of NLS prediction programs, we analyzed the localization of GFP-Piwi6 containing five sequential deletions in the IDR (size range 19 to 28 amino acids; Supplementary Figure S8A). Deletion of residues 29–56 (Δ29-56), but none of the other deletions, strongly reduced nuclear localization (Supplementary Figure S8B-C). Interestingly, the Δ29-56 mutant and the ΔNLS2 mutant (residues 52–85) overlap in a short sequence patch of charged amino acids (aa 48-61: RGDHRQKPYDRPEH), conserved in *Ae. aegypti* Piwi6 but absent from Piwi5 (Supplementary Figure S1), which we propose may be the main determinant of Piwi6 nuclear localization.

**Figure 5.**
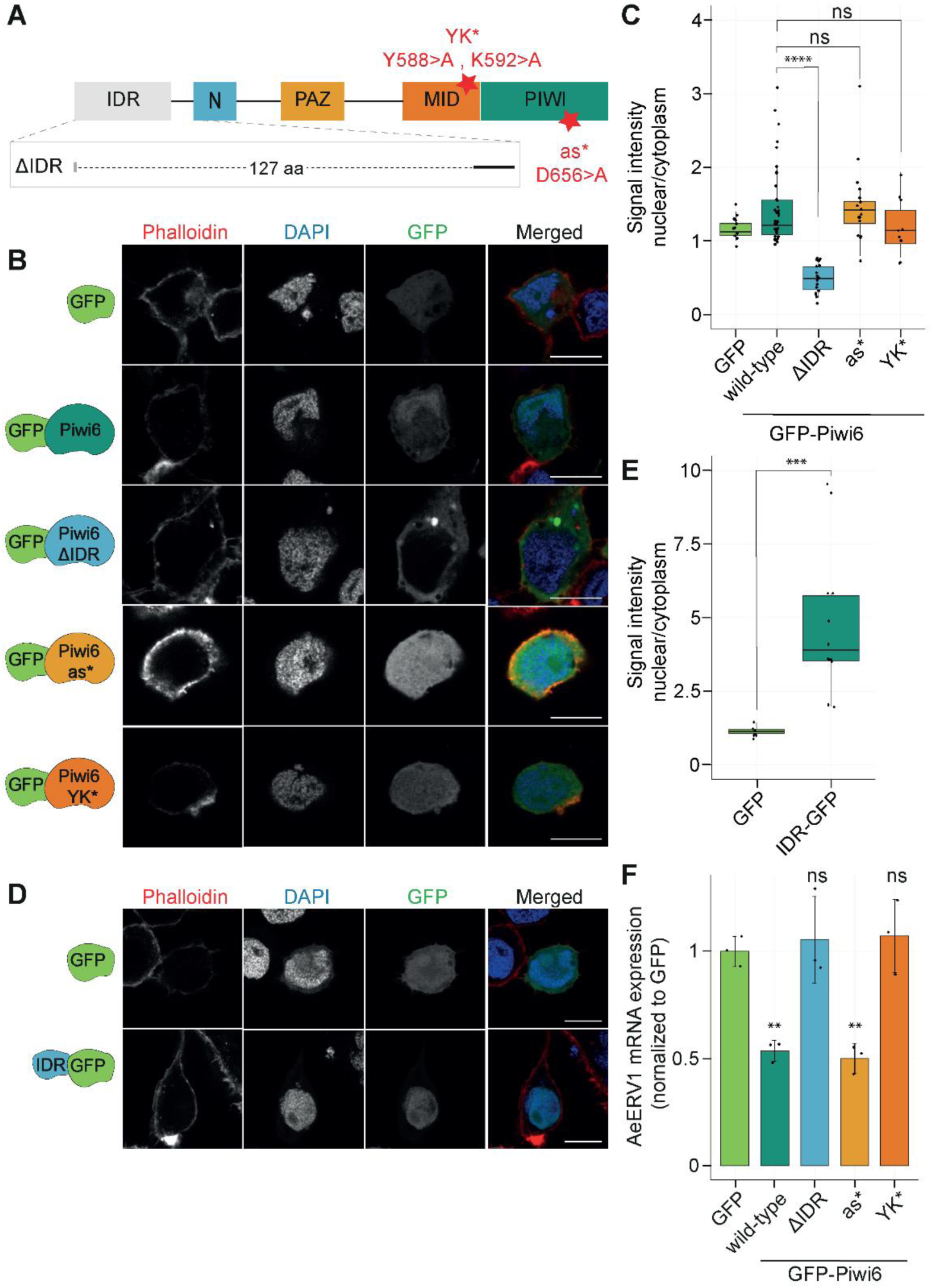
Piwi6 nuclear localization is required for *AalERV1* silencing. A) Domain structure of *Ae. albopictus* Piwi6. The deletion of the intrinsically disordered region (ΔIDR) is indicated with dashed lines, and mutated residues in the putative active site (as*) and the 5’ piRNA recognition site (YK*) are shown in red. The domain structure was based on a sequence alignment with *B. mori* Siwi, using the published crystal structure [61]. B) Representative confocal images showing the subcellular localization of GFP, GFP-Piwi6, and the indicated mutants in *Piwi6* KO cells. C) Quantification of the GFP signal intensity from panel B, presented as the ratio of nuclear to cytoplasmic expression. The centerline in the boxplots represents the mean, box edges indicate the first and third quartiles, whiskers show maximum and minimum values. Statistical significance was tested against GFP-tagged wild-type Piwi6. Dots represent individual cells: GFP (n = 12), wild-type *Piwi6* (n = 41), ΔIDR (n = 20), AS* (n = 16), and YK* (n = 10). D) Representative confocal images of GFP and a PIWI6 IDR-GFP fusion protein in *Piwi6* KO cells. E) Quantification of the GFP signal of panel (E), showing the ratio of the nuclear to cytoplasmic GFP signal intensity. Dots represent individual cells: GFP (n = 8) and IDR-GFP (n = 12). Statistical significance was tested against the GFP control. One-way ANOVA with Holm-Sidak’s multiple comparisons post hoc test was performed in B and D, and two-tailed student’s *t*-tests were performed in F to test the statistical significance; ** *P* < 0.01, *** *P* < 0.001, **** *P* < 0.0001, ns, not significant. In C and E, DAPI and phalloidin staining were used to stain the nuclei and cytoplasm, respectively. Uncropped images are shown in Supplementary Figure S10. Scale bars represent 10 μm. F) *AalERV1* mRNA expression in *Piwi6* KO cells expressing a GFP-tagged wild-type *Piwi6* transgene or the indicated mutants, measured by RT-qPCR. *AalERV1* expression was normalized to the house-keeping gene *RPL5* and expressed relative to *Piwi6* KO cells expressing the GFP control. Bars represent the mean ± SD of three biological replicates in an experiment representative of two independent experiments. Statistical significance was tested against the GFP control. See also Supplementary Figure S7 and S10.

### The catalytic tetrad is not required for *AalERV1* silencing

To correlate the subcellular localization of Piwi6 with its function in *AalERV1* silencing, we performed genetic rescue experiments by expressing the wild-type or mutant *GFP-Piwi6* transgene in *Piwi6* KO cells. GFP-Piwi6 was expressed at levels comparable to endogenous Piwi6 at two days after transfection (Supplementary Figure S7F), and *AalERV1* mRNA levels were significantly reduced in cells expressing GFP-Piwi6 compared to cells expressing GFP as a control (Figure 5F), indicating that the fusion protein mediates functional silencing. We hypothesized that loss of the nuclear localization of Piwi6 would lead to a defect in *AalERV1* silencing. Indeed, GFP-Piwi6-ΔIDR failed to rescue the defect in *AalERV1* silencing of *Piwi6* KO cells, in stark contrast to wild-type GFP-Piwi6 (Figure 5F). The catalytic tetrad mutant (as*) rescued *AalERV1* silencing to a similar extent as wild-type GFP-Piwi6 (Figure 5F), indicating that *AalERV1* silencing is independent of a putative slicer activity of Piwi6. As anticipated for a mutant unable to bind piRNAs [9], the GFP-Piwi6-YK* mutant did not rescue *AalERV1* silencing (Figure 5F and Supplementary Figure S7A). However, failure to rescue silencing may also be due to reduced GFP-Piwi6-YK* protein expression (Supplementary Figure S7G), suggesting that unloaded Piwi6 is unstable (Supplementary Figure S7A), as previously observed for other PIWI proteins [72,79,80].

### Piwi6 silences *AalERV1* both in ovaries and somatic tissues

U4.4 cells are derived from *Ae. albopictus* larvae, and their PIWI protein and piRNA expression patterns resemble those of somatic tissues [29]. To assess the role of Piwi6 in *AalERV1* silencing *in vivo*, we performed intrathoracic injections of dsRNA targeting *Piwi6* in female mosquitoes and evaluated *AalERV1* expression in ovaries and the rest of the carcass (somatic tissues) (Figure 6A). After injection, the mosquitoes were allowed to blood feed to initiate ovary development. Despite a relatively modest knockdown efficiency of about 50% (Figure 6B), *AalERV1* expression was significantly increased 6-fold in carcasses and 2-fold in ovaries relative to mosquitoes treated with control dsRNA (Figure 6C). The stronger effect in carcasses could not be explained by differences in knockdown efficiencies, suggesting that Piwi6 is particularly active in the soma. Within the same samples, *Piwi4* and *Piwi5* expression were not significantly affected by *Piwi6* knockdown (Supplementary Figure S9), validating the knockdown specificity. These experiments confirm our observations in cell lines and highlight the importance of the somatic piRNA pathway, and specifically Piwi6, in TE silencing *in vivo*.

**Figure 6.**
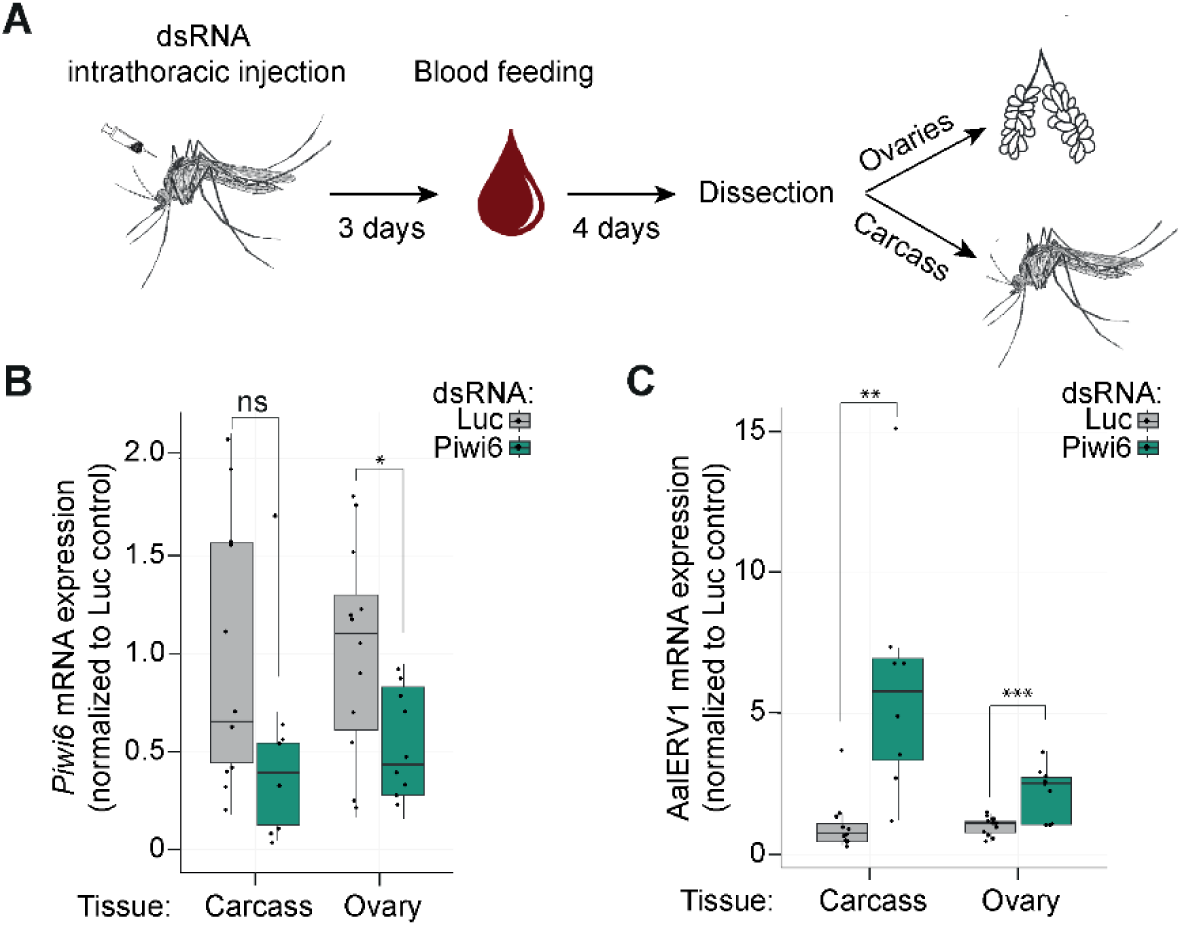
*In vivo Piwi6* knockdown results in transposon upregulation in ovaries and somatic tissues. A) Outline of the *in vivo* knockdown experiment. B–C) *Piwi6* (B) and *AalERV1* expression (C) in ovaries and carcasses from mosquitoes injected with dsRNA targeting *Piwi6* or firefly *luciferase* (Luc), assessed by RT-qPCR. mRNA expression of target genes was normalized to the housekeeping gene *RPL5* and expressed relative to the non-targeting control dsRNA (Luc). The centerline in the boxplots shows the mean, box edges represent the first and the third quartile, and the whiskers show maximum and minimum values. Two-tailed student’s *t*-tests were used to determine statistically significant differences with control (Luc), * *P* < 0.05; ** *P* < 0.01; *** *P* < 0.001, ns, not significant. Dots represent individual samples analyzed in carcass: Luc (n = 11) and Piwi6 (n = 8) and in ovaries: Luc (n = 12) and Piwi6 (n = 9). See also Supplementary Figure S9.

## Discussion

Since its discovery in *Drosophila*, the piRNA pathway has been thought to be a gonad-specific defense mechanism against TE invasion. This notion was challenged by comparative analyses of diverse arthropod species, suggesting that non-gonadal piRNA expression is an evolutionary conserved trait of the arthropod phylum [13], which was lost in *Drosophila*. *Aedes* mosquitoes are among the organisms in which the piRNA pathway is active in both somatic tissues and the gonads. Of particular interest, the closely related *Ae. albopictus* and *Ae. aegypti* encode an expanded PIWI gene family, of which four members, Piwi4–6 and Ago3, are expressed in somatic tissues. Using *Piwi6* knockouts in the *Ae. albopictus* U4.4 cell line, we here show that Piwi6 mediates transcriptional gene silencing of TEs, specifically the class of LTR retrotransposons. Transcriptional silencing was associated with the deposition of the repressive heterochromatin mark H3K9me3 and relied on the nuclear localization but not the catalytic tetrad of Piwi6. Furthermore, we identified a full-length endogenous retrovirus, named *AalERV1*, that was particularly sensitive to Piwi6-mediated silencing, and we speculate that transcriptional silencing preferentially targets active elements. Finally, we found that *AalERV1* is silenced by the piRNA machinery in somatic tissues, establishing *Ae. albopictus* as a novel model to study nuclear piRNA mechanisms.

Our data supports the following model. In wildtype conditions, Piwi6 is loaded with antisense piRNAs targeting TEs including *AalERV1*. Due to the NLS in its intrinsically disordered region, Piwi6 translocates to the nucleus to recognize nascent target transcripts and recruit histone modifying enzymes to deposit the repressive H3K9me3 mark. Loss of the repressive chromatin structure upon *Piwi6* knockout leads to enhanced RNA polymerase II occupancy and increased transcription. The resulting transcripts may then be targeted by Piwi5 loaded with antisense piRNAs expressed from piRNA clusters, which may then engage in ping-pong amplification with Ago3, explaining the observed increase numbers of both sense and antisense piRNAs. In *Drosophila*, PIWI protein knockout leads to loss of the piRNA population that it binds [10,63,81]. We propose that unaffected *Piwi5* expression explains why piRNAs levels do not collapse in *Piwi6* KO cells, as both Piwi5 and Piwi6 bind antisense TE-derived piRNAs [25,29]. It is possible that Piwi6 only receives a small fraction of primary, cluster-derived piRNAs, explaining why its loss does not significantly affect the global piRNA production.

### A somatic piRNA pathway mediates TE silencing

Our data suggest that Piwi6 plays a crucial role in regulating young and potentially active LTR retrotransposons in the soma. While *Piwi6* knockdown derepressed *AalERV1* expression in somatic tissues as well as in the ovary, our data do not allow us to discriminate whether Piwi6 is active in the germline or the somatic cells of the ovary. Our results therefore support the recent proposal that the somatic piRNA pathway is an ancestral TE defense mechanism across arthropods [13]. Moreover, our findings offer an opportunity to study how endogenous retroviruses are transmitted and expand in the genome [82]. For example, it will be of interest to study whether *AalERV1* forms viral particles in the soma to invade the germline, akin to mobile elements such as Gypsy and ZAM in *Drosophila melanogaster* [83–85].

### A non-model organism to study nuclear piRNA pathway

Small RNA-guided heterochromatin formation is remarkably conserved across eukaryotes. Organisms ranging from fission yeast to mammals use small RNAs to target specific loci for heterochromatin formation [86]. Yet, although the core concepts of small RNA-guided heterochromatin formation are conserved, the mechanisms differ greatly. In the fission yeast *Schizosaccharomyces pombe*, small interfering (si)RNA-loaded Ago1 silences centromeric repeat derived transcripts through H3K9me3 deposition and initiation of heterochromatin formation [87,88]. In mice, the nuclear PIWI protein MIWI2 triggers *de novo* DNA methylation of target retrotransposons during male germ cell development [89,90]. In *Arabidopsis thaliana*, heterochromatic siRNAs drive Argonaute 4 into the nucleus to silence targets via DNA methylation [91].

The mosquito piRNA pathway holds great potential for studying nuclear PIWI protein function and evolution of central components of the transcriptional gene silencing machinery. In *Drosophila*, transcriptional gene silencing depends on the recruitment of multiple factors to target loci by the Piwi-piRNA complex, such as the nuclear Panoramix-induced co-transcriptional silencing complex (PICTS/SFiNX), with Panoramix (Panx) and its cofactor Cut-up as its primary components [92,93]. *Aedes* mosquitoes encode a Cut-up ortholog and a putative ortholog of *Panx* that is highly divergent from *Drosophila* [92], suggesting rapid evolution of this gene under positive selection. Additionally, *Aedes* mosquitoes encode other PICTS components, Nxf1 and Nxt1, as well as cofactors such as Maelstrom and Asterix; however, whether similar protein complexes are required for transcriptional silencing in mosquitoes remains unknown.

### The intrinsically disordered region is essential for Piwi6 function

We found that the Piwi6 IDR was essential for nuclear localization and transposon silencing and that it functions as a NLS when fused to the N-terminus of GFP. As NLSs are typically short, positively charged peptides [94], the 127 aa long IDR is unlikely to be the NLS itself. Sequences spanning residues residues 29–56 and residues 52–85 within the IDR contributed to Piwi6 nuclear localization. Notably, the Piwi6 IDR sequence does not seem to match known NLS categories, including monopartite or bipartite, or non-canonical NLSs [94], and the specific importin proteins facilitating Piwi6’s nuclear transport remain unknown. Moreover, our rescue assay using a piRNA-binding defective mutant indicate that piRNA loading is not essential for Piwi6 nuclear localization, contrasting with the bipartite canonical NLS of *Drosophila* Piwi, which becomes exposed to Impα upon piRNA loading due to a conformational change [73].

### Transcriptional silencing does not require the catalytic tetrad

*Drosophila* Piwi lacks slicer activity due to its non-canonical DVDK tetrad, and it has been proposed that this prolongs association with target RNA, allowing Piwi to serve as an RNA binding platform [76]. Although Piwi6 contains the canonical DEDH catalytic tetrad, slicer activity has not been shown in biochemical assays and it is possible that residues involved in the hydrogen bonding network around the catalytic tetrad and other structural constraints may prevent slicer activity or render it very inefficient [61]. Interestingly, our rescue assays suggest that the catalytic tetrad is not required for transposon silencing by Piwi6, in line with its function in transcriptional silencing in the nucleus. Yet, Piwi6 is also expressed in the cytoplasm, and we cannot rule out that it participates in post transcriptional silencing as well, possibly via interactions with other cofactors in the cytoplasm.

### *Is AalERV1* is a mobile element*?*

Despite the high number of TE insertions, piRNA clusters in *Aedes* genomes are depleted of most TE sequences relative to the entire genome, with the exception of LTR retrotransposon and rolling-circle DNA transposons, which are slightly enriched [40]. This is in stark contrast to *Drosophila* where piRNA clusters are highly enriched in TE fragments. Interestingly, several copies of *AalERV1* seem to encode complete *gag*, *pol* and *env* genes [67] flanked by identical LTRs, and we speculate that *AalERV1* is an active element that has transposed recently. By aligning the consensus *AalERV1* sequence against the *Ae. albopictus* genome [40] using BLAST, we identified seven insertions in four distinct piRNA clusters (Supplementary Table S11). Two of these clusters contained nearly the entire *AalERV1* sequence, either as a single insertion with gaps (cluster 96, nt identity > 97%) or as multiple insertions collectively covering the sequence (cluster 53, nt identity > 94%). The two remaining clusters, categorized as core clusters in [29], contained partial insertions of the 3’ end of *AalERV1* sequence (cluster 2 and 18, nt identity > 88%). Together, these analyses suggest that full-length elements are kept in check through transcriptional silencing via cluster derived piRNAs. However, increased *AalERV1* mRNA expression does not guarantee mobilization within the genome [95] and whether *AalERV1* is transposition competent remains to be demonstrated.

In summary, our study uncovers the pivotal role of Piwi6 in transcriptional silencing of TEs in the soma of mosquitoes. We propose that Piwi6 is particularly active against LTR retrotransposons, including young, full-length endogenous retroviruses. Transcriptional silencing was independent of the catalytic triad but required a nuclear localization signal in the IDR of Piwi6. Our results establish *Aedes* mosquitoes as a model to study nuclear PIWI function and suggest that somatic piRNA-mediated transposon silencing is evolutionarily conserved across arthropods.

## Data availability

Small RNA-seq and RNA-seq data have been deposited in NCBI Sequence Read Archive under accession numbers PRJNA989754; PRJNA1173503; PRJNA989754; PRJNA1173503; PRJNA1173503. PRO-seq and CUT&Tag data, as well as small RNA-seq data from GFP-IPs have been deposited in NCBI Sequence Read Archive under accession numbers PRJNA1470531. Other data underlying this article are available in the article and in its online supplementary material. Scripts used in bioinformatics analyses will be deposited in GitHub and become available upon publication.

## Acknowledgements

We thank past and current members of the laboratory for discussions. *Ae. albopictus* mosquitoes were kindly provided by Fabrice Chandre and Bethsabée Scheid (Institut de Recherche pour le Développement, Montpellier) through the InfraVec2 program. We thank the Radboudumc Technology Center Microscopy for use of their microscopy facilities.

## Author Contributions

Conceptualization, E.T. and R.P.v.R.; Methodology, E.T., N.B.v.E., F.A.H.v.H.; M.B., G.J.O, and R.P.v.R; Formal Analysis, E.T., N.B.v.E., F.A.H.v.H.; Investigation, E.T., N.B.v.E., F.A.H.v.H., M.B., G.J.O., C.L., R.H.; Writing – Original Draft, E.T. and R.P.v.R.; Writing – Review & Editing, E.T., N.B.v.E., F.A.H.v.H.; M.B., G.J.O., C.L., C.J.M.K., J.Q., P.M., R.H and R.P.v.R.; Visualization, E.T., N.B.v.E., G.J.O.; Supervision, C.J.M.K., P.M., R.H. and R.P.v.R.; Funding Acquisition, R.P.v.R.

