## Supplementary Figures 1-10, Supplementary Tables 1-11 for "The piRNA pathway mediates transcriptional silencing of LTR retrotransposons in ovaries and somatic tissues of *Aedes* mosquitoes"

S1

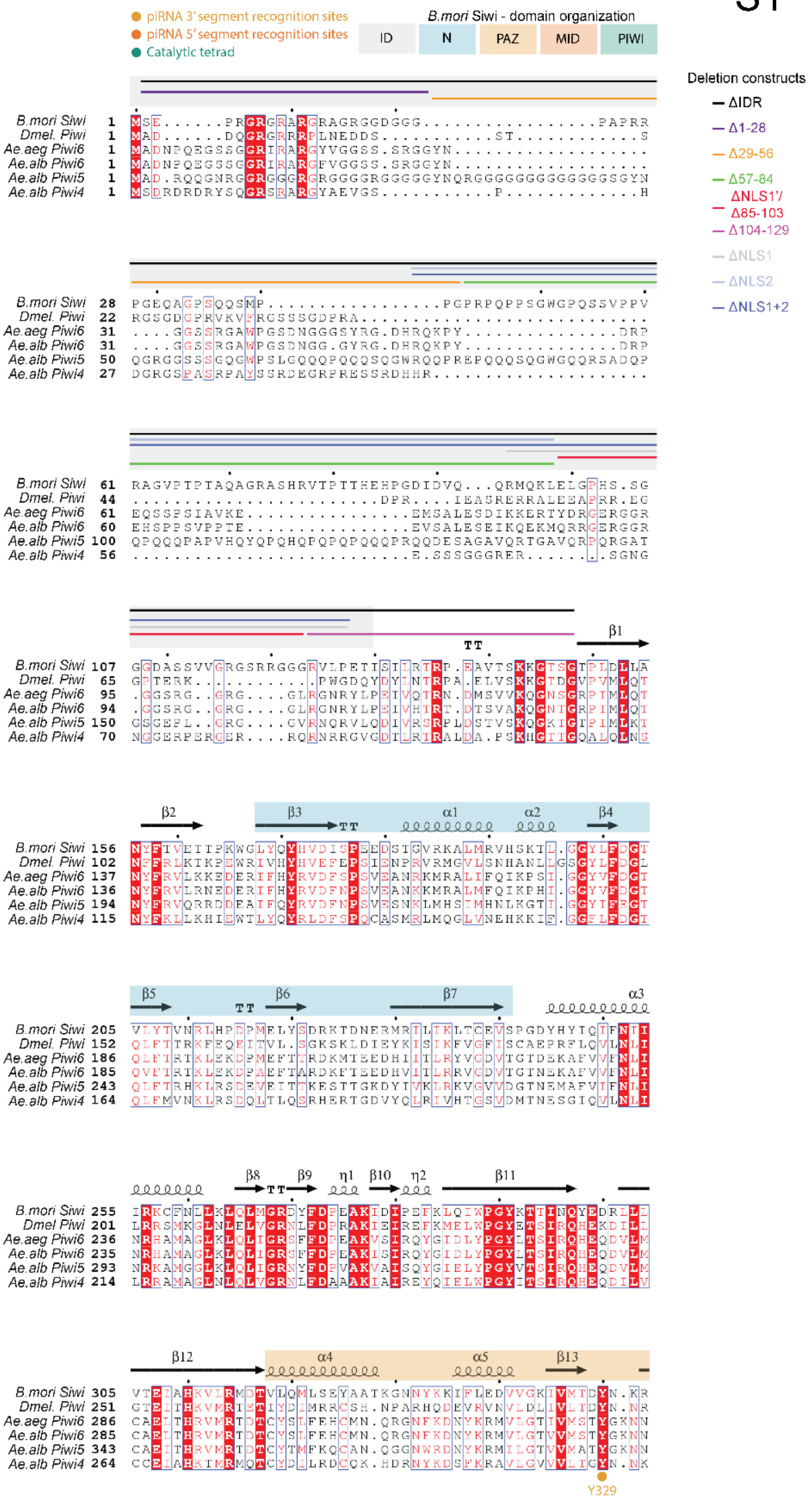
piRNA 3' segment recognition sites
piRNA 5' segment recognition sites
Catalytic tetrad
B.mori Siwi - domain organization
ID
N
PAZ
MID
PIWI
Deletion constructs
ΔIDR
Δ1-28
Δ29-56
Δ57-84
ΔNLS1'/ Δ85-103
Δ104-129
ΔNLS1
ΔNLS2
ΔNLS1+2
B.mori Siwi
Dmel. Piwi
Ae.aeg Piwi6
Ae.alb Piwi6
Ae.alb Piwi5
Ae.alb Piwi4
Y329

S1

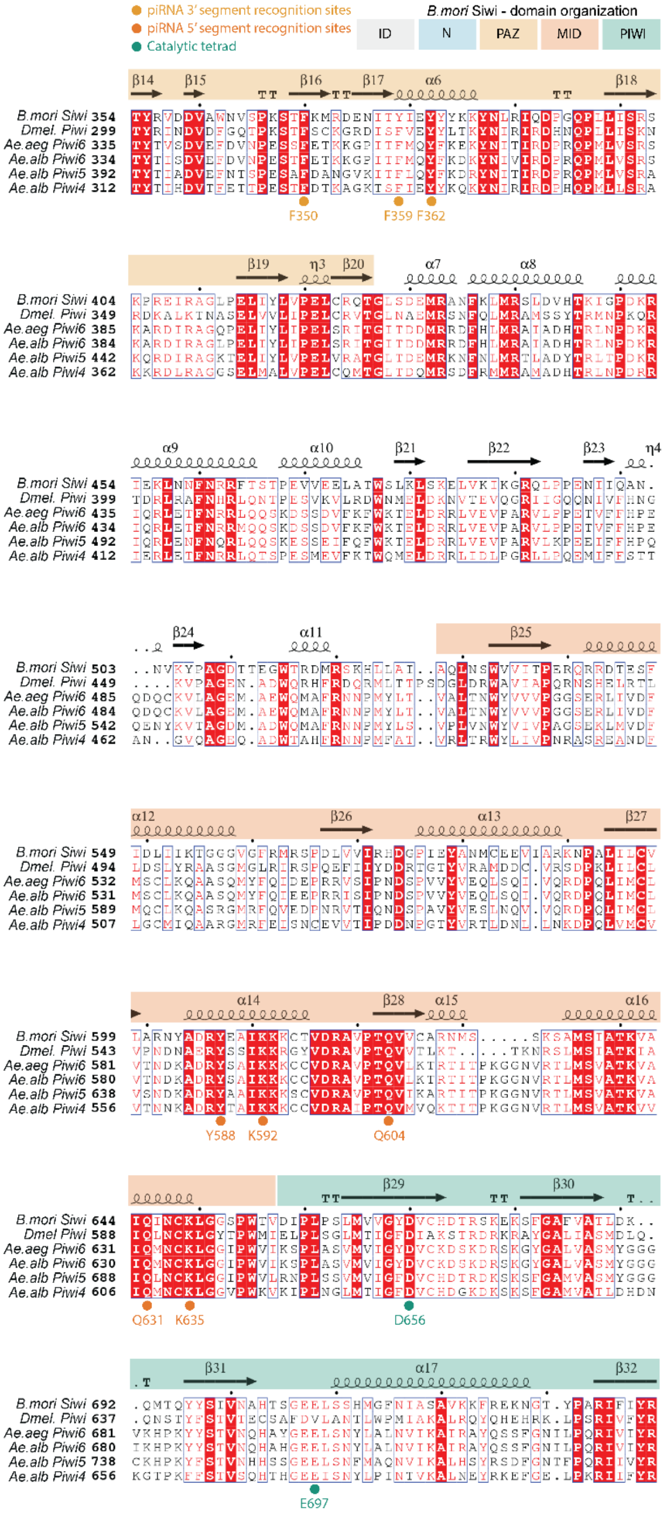
piRNA 3' segment recognition sites
piRNA 5' segment recognition sites
Catalytic tetrad
B.mori Siwi - domain organization
ID
N
PAZ
MID
PIWI
B.mori Siwi
Dmel. Piwi
Ae.aeg Piwi6
Ae.alb Piwi6
Ae.alb Piwi5
Ae.alb Piwi4
F350
F359 F362
Y588 K592
Q604
Q631 K635
D656
E697

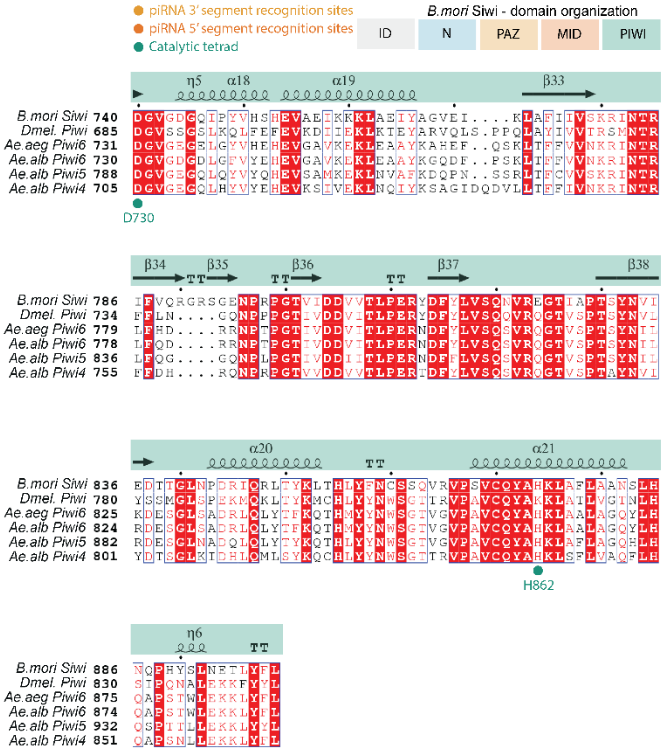


**Figure S1. Multiple sequence alignment of PIWI proteins.**

Multiple sequence alignment of *B. mori* Siwi, *D. melanogaster* Piwi, *Ae. albopictus* Piwi4 and Piwi5, and *Ae. aegypti* and *Ae. albopictus* Piwi6. The secondary structure of Siwi is indicated above the sequence alignments [61]. Different domains of *Ae. albopictus* Piwi6 are indicated with different colors. Crucial sites are indicated with dots, and their corresponding positions in *Ae. albopictus* Piwi6 are indicated below the sequence alignments. The position of deletion mutants are indicated with horizontal lines.

S2

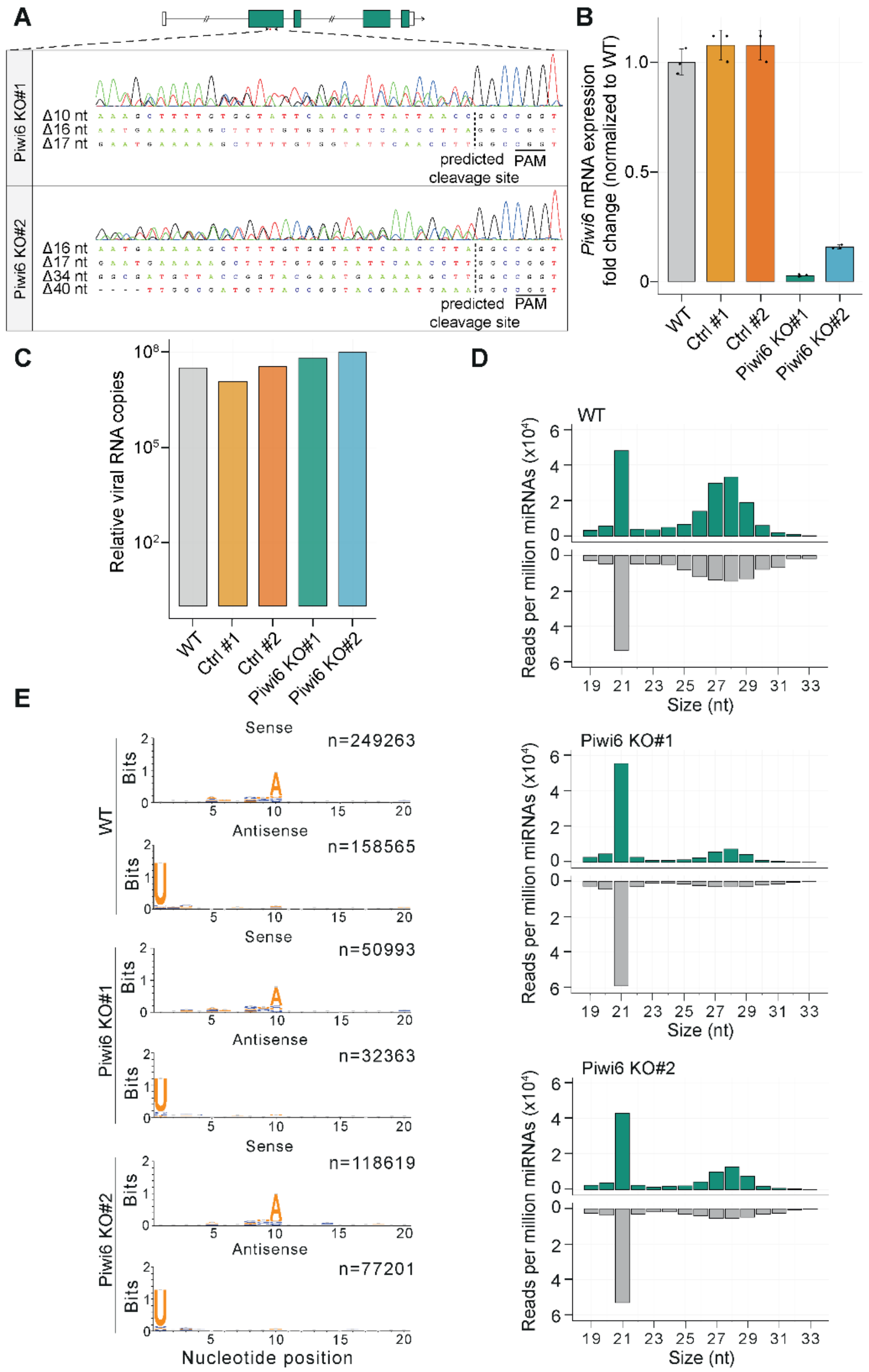
A
Piwi6 KO#1
Δ10 nt
Δ16 nt
Δ17 nt
predicted cleavage site
PAM
Piwi6 KO#2
Δ16 nt
Δ17 nt
Δ34 nt
Δ40 nt
B
Piwi6 mRNA expression fold change (normalized to WT)
WT
Ctrl #1
Ctrl #2
Piwi6 KO#1
Piwi6 KO#2
C
Relative viral RNA copies
D
WT
Piwi6 KO#1
Piwi6 KO#2
Reads per million miRNAs (x10⁴)
Size (nt)
E
Sense
Antisense
n=249263
n=158565
n=50993
n=32363
n=118619
n=77201
Bits
Nucleotide position

**Figure S2. Validation of *Piwi6* knockout in *Ae. albopictus* U4.4 cells.**

A) *Piwi6* KO cells were generated by CRISPR/Cas9-mediated editing of the *Piwi6* gene in U4.4 cells. Top panel shows the gene structure of *Piwi6* (*AALFPA_065618*) annotated in the *Ae. albopictus* genome Aalbo_primary1 (RefSeq GCF_006496715.1). Bottom panel: Sanger sequencing results of two *Piwi6* KO cells (Piwi6 KO#1: guide #1, clone #10 and Piwi6 KO#2: guide #1, clone #52) containing only out-of-frame deletions. The predicted cleavage sites upstream of the PAM sequence are indicated by dashed vertical lines.

B) *Piwi6* expression in WT, control (Ctrl) and *Piwi6* KO cells, assessed by RT-qPCR. mRNA expression was normalized to WT samples. The housekeeping gene *RPL5* was used as an internal control. Bars indicate mean ± SD of three biological replicates in an experiment representative of two independent experiments.

C) Relative Sindbis virus RNA copies in WT, Ctrl and *Piwi6* KO cells, assessed by RT-qPCR using primers for the *NSP4* gene (n = 1, the same samples were used for deep sequencing shown in panel D).

D) Read size distribution of 19–33 nt RNA reads mapping to the sense (green) and antisense (grey) strands of the Sindbis virus (SINV-GFP) genome, allowing one mismatch in WT (top panel), *Piwi6* KO#1 (middle panel) and *Piwi6* KO#2 (bottom panel) cells. Reads were normalized to the number of miRNAs (in millions).

E) Sequence logos of the first 20 nucleotides of piRNAs mapping to the SINV-GFP genome in sense and antisense orientation in WT (top panel), *Piwi6* KO#1 (middle panel) and *Piwi6* KO#2 (bottom panel) cells. Number of reads contributing to sequence logos are indicated at the top right of each sequence logo.

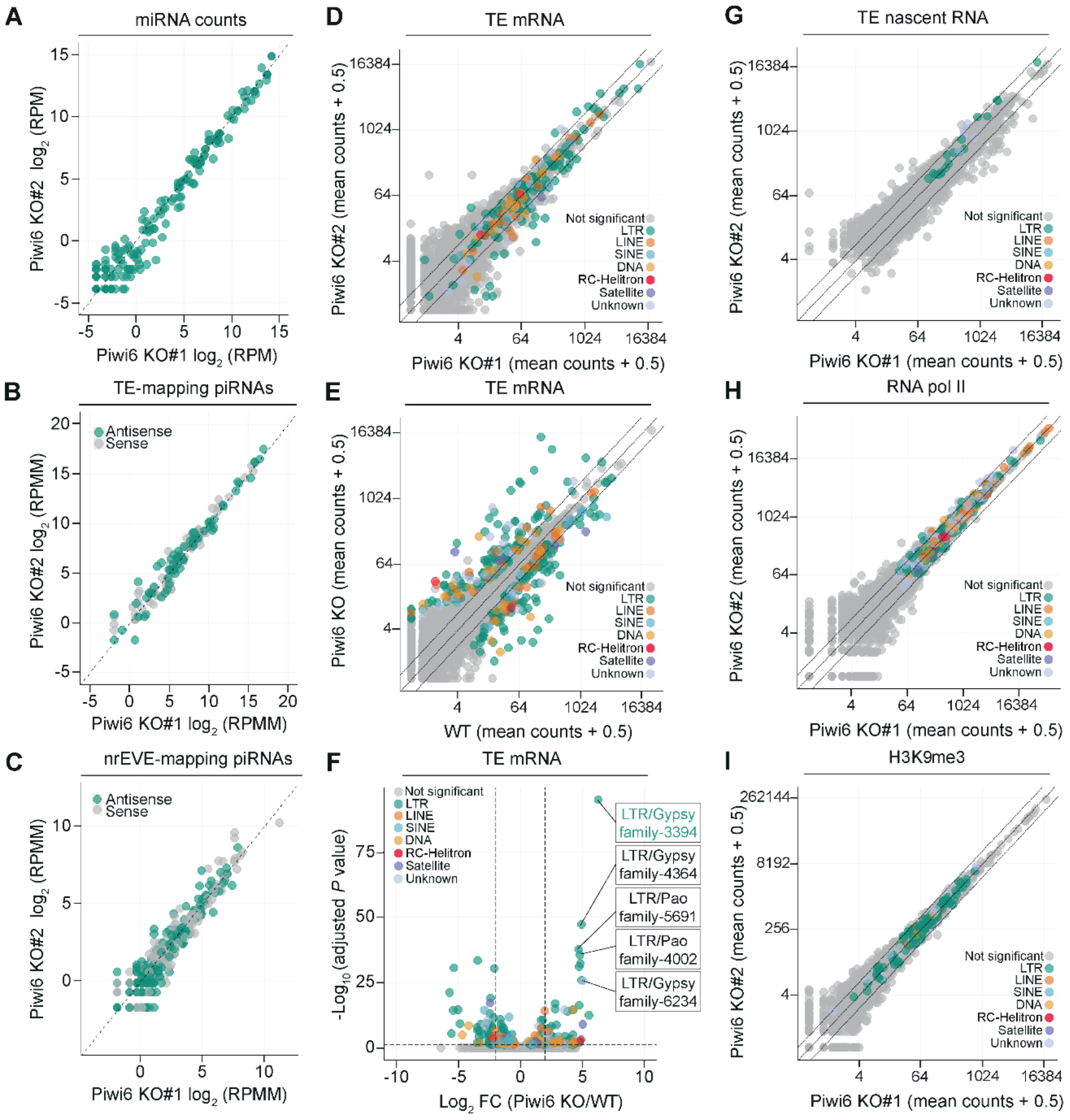


**Figure S3. Correlation of small RNA, TE, and gene expression between the two *Piwi6* KO clones.**
A) Scatterplots showing miRNA expression in two *Piwi6* KO clones. Reads were counted, one pseudo count was added, and read counts were normalized to library size (RPM, reads per million).
B-C) Scatterplots of piRNAs mapping to TEs (B) and nrEVEs (C) in two *Piwi6* KO clones. One pseudo count was added to the raw counts, and the reads were normalized to miRNAs (in millions) (RPMM).
D) mRNA expression of TEs in two *Piwi6* KO clones. Statistical testing was performed as described in Figure 3A for TEs and statistical significance of differential expression was tested against control clones (Ctrl). Gray symbols, not significant; colored symbols, differentially expressed TEs, categorized according to the *Ae. albopictus* repeat annotation [41].
E) Expression of TE mRNAs in WT and *Piwi6* KO cells measured by RNA-seq. The same analysis as in Figure 3A was performed with statistical significance of differential expression was tested against WT cells instead of control clones (Ctrl).

F) Volcano plot showing the $\log_2$ fold changes in TE mRNA expression between *Piwi6* KO cells and WT cells. The same analysis as in Figure 3B was performed with statistical significance of differential expression was tested against WT cells instead of control clones (Ctrl).

G-I) Scatterplots of nascent TE RNA levels (G), RNA polymerase II occupancy (RNA pol II; H) and H3K9me3 enrichment (I) at TE loci in two *Piwi6* KO clones. Reads were normalized using DESeq2, with a pseudo count of 0.5 added. Gray symbols, not significant; colored symbols, differentially expressed TEs categorized according to the *Ae. Albopictus* repeat annotation [41], modified to replace LTR/Gypsy family 3394 with the *AalERV1* consensus sequence.

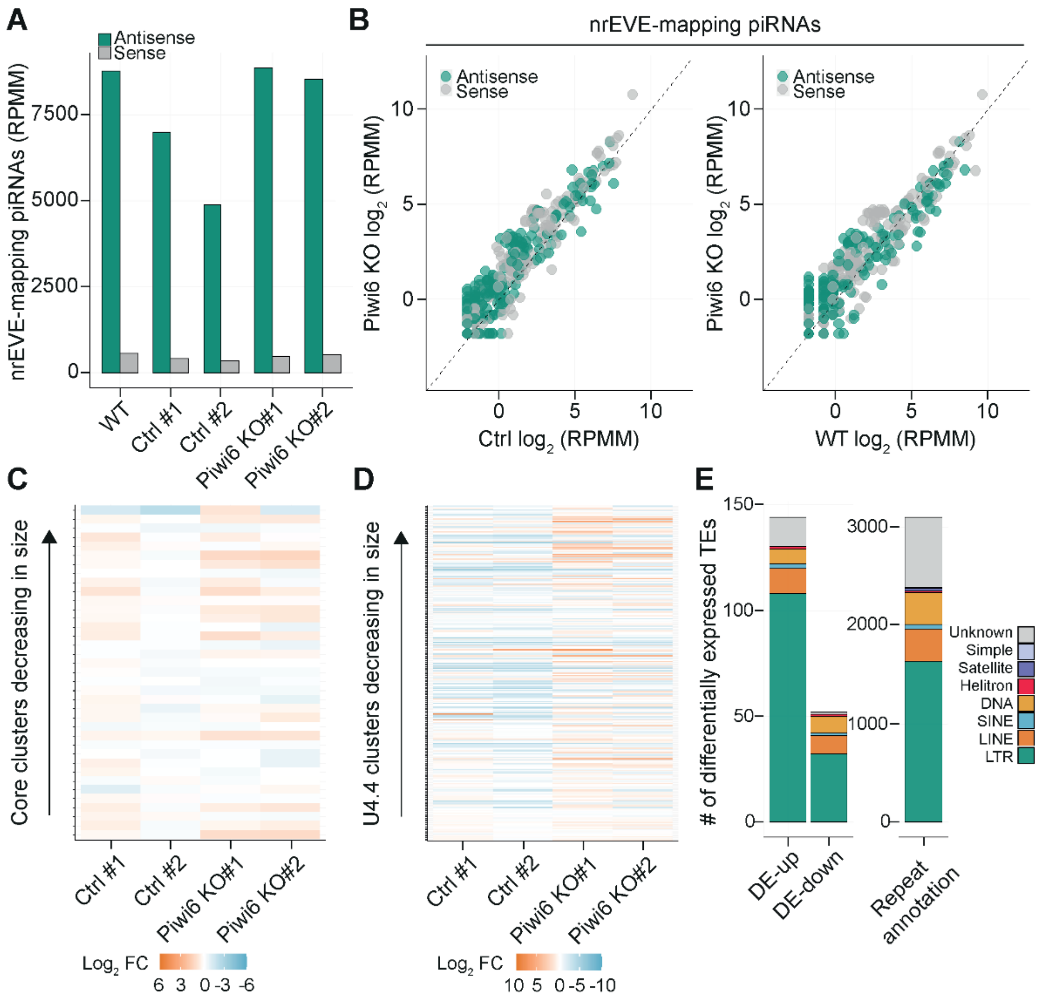


**Figure S4. Endogenous piRNA and TE expression in *Ae. albopictus Piwi6* knockout cells.**

A) Bar plots showing the number of piRNAs mapping to non-retroviral endogenous viral elements (nrEVEs) in sense (gray) and antisense (green) orientation in WT, Ctrl and *Piwi6* KO cells. Reads were normalized to the number of miRNAs (in millions) and expressed as reads per million miRNAs (RPMM).

B) Scatterplots comparing piRNAs mapping to nrEVEs in WT, Ctrl and *Piwi6* KO cells. nrEVE-mapping reads were counted, a pseudo count of one was added, counts were normalized to the number of miRNAs (in millions) and expressed as $log_2$-transformed reads per million miRNAs (RPMM). For Ctrl and *Piwi6* KO clones the mean of two clones is shown. Sense (gray) and antisense (green) mapping strand was determined relative to the orientation of predicted viral open reading frames of nrEVEs.

C-D) Heatmap comparing piRNA expression in Ctrl and *Piwi6* KO to WT cells, mapping to core piRNA clusters (C) and all U4.4 clusters (D) defined in [29]. Reads mapping to the clusters were counted, normalized to the number of miRNAs (in millions) and displayed as $log_2$ fold change (FC) in a heatmap. Clusters were ranked by size.

E) Stacked bar graphs indicating the number of differentially expressed (DE) TEs between *Piwi6* KO cells and Ctrl cells (DE-up, upregulated; DE-down, downregulated) categorized according to TE families *Ae. albopictus* repeat annotation [41]. The rightmost bar shows the number of TE families in the *Ae. albopictus* repeat annotation [41].

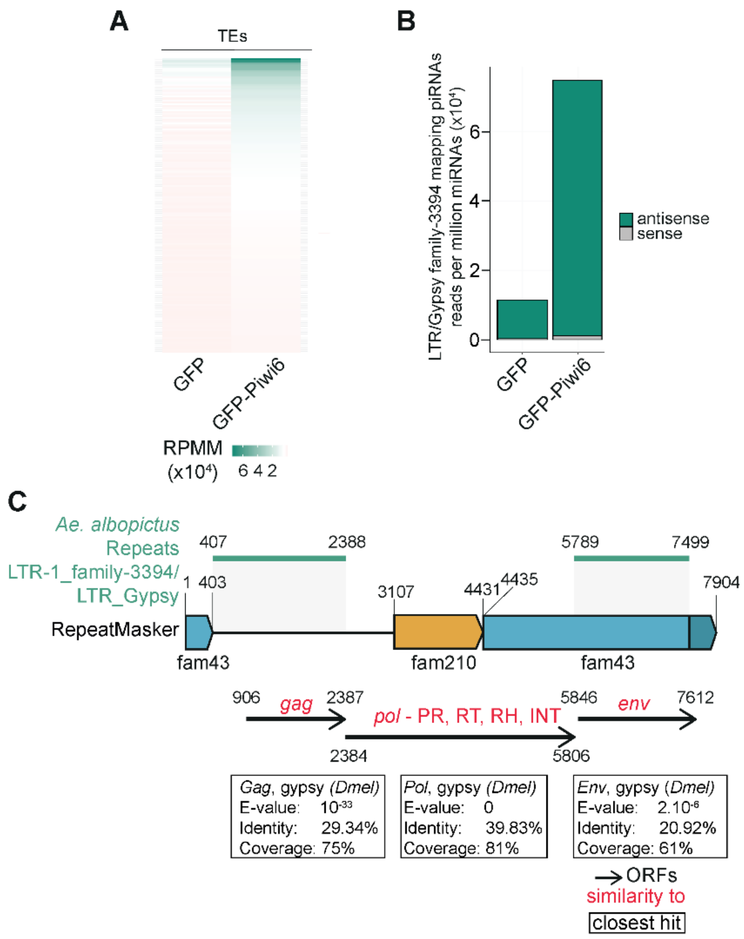


**Figure S5. Piwi6 associated piRNAs and schematic representation of *AalERV1*.**

A) Heatmap showing the TE-derived piRNAs enriched in GFP-tagged Piwi6 immunoprecipitation compared to the GFP control. Small RNA reads were mapped to *Ae. albopictus* repeat annotation [41] allowing one mismatch, read counts were normalized to the total number of miRNAs (in millions) (RPMM). TE elements showing > 2-fold enrichment in GFP-Piwi6 over GFP and normalized counts > 1000 RPMM were included. TEs are ranked by read counts.

B) piRNA read counts mapping to LTR Gypsy-family 3394 element in antisense and sense orientation in GFP-tagged Piwi6 and GFP immunoprecipitation. Small RNA reads were mapped to *Ae. albopictus* repeat annotation allowing one mismatch, read counts were normalized to the total number of miRNAs (in millions) (RPMM).

C) TE elements in the indicated *AalERV1* locus annotated in the *Ae. albopictus* genome by RepeatMasker [40] are highlighted as colored boxes. Identical 5' LTR and 3' LTR are shown in dark blue. Gray shading indicates the alignment of the LTR Gypsy-family 3394 element (in green) of the *Ae. albopictus* repeat annotation [41] to the indicated *AalERV1* locus. Open reading frames predicted by ORFfinder [50] using the entire genomic sequence are highlighted with black arrows. Similarities of the ORF sequences to UniProtKB/Swiss-Prot database are indicated in red. PR, protease; RT, reverse transcriptase; RH, RNase H; INT, integrase; env, envelope. ORFs are shown in Supplementary Table S9.

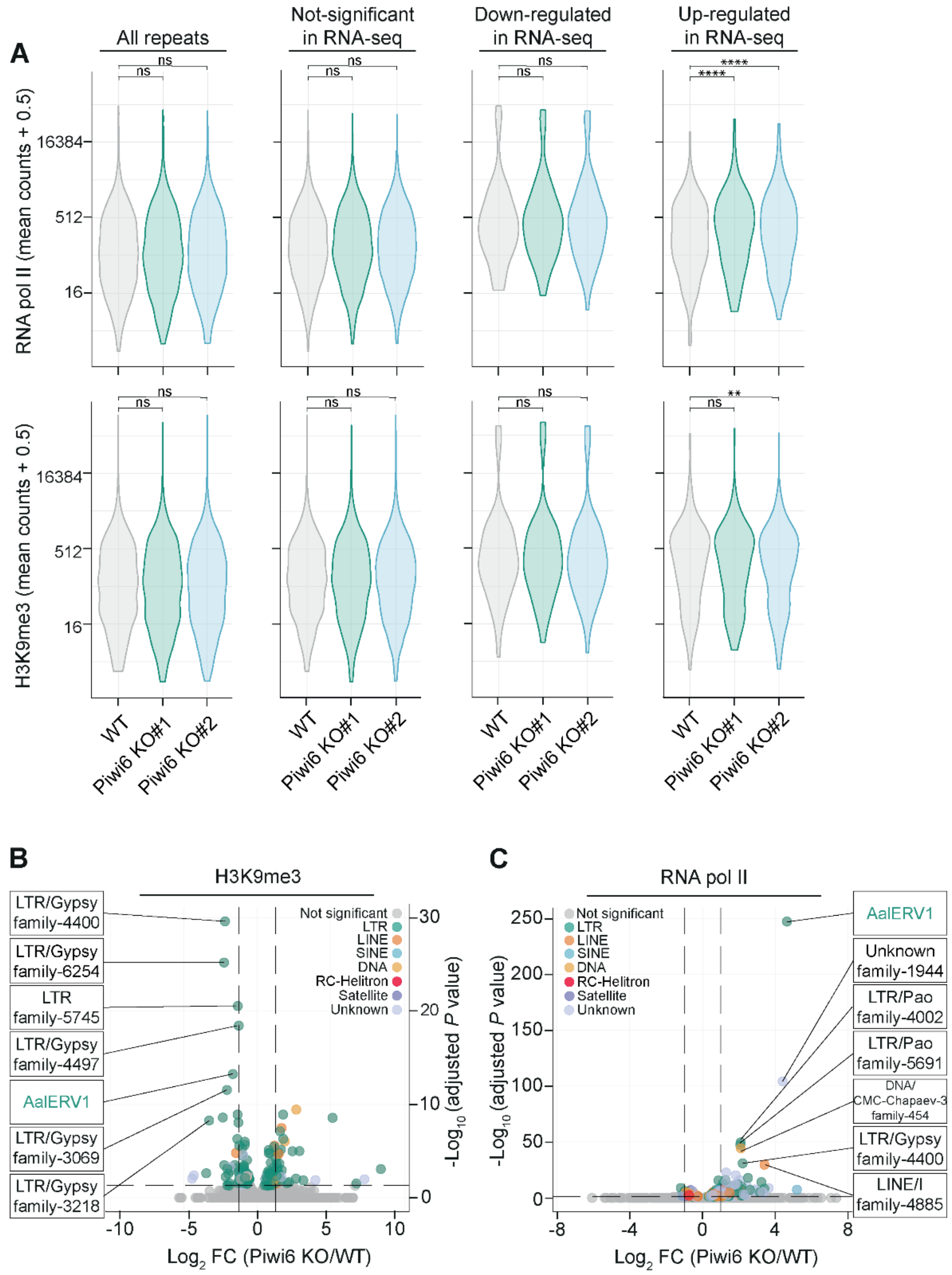


**Figure S6. RNA pol II occupancy and H3K9me3 enrichment at TE loci**

A) Enrichment of RNA polymerase II (RNA pol II) and H3K9me3 at TE loci in WT and *Piwi6* KO cells, as measured by CUT&Tag. Reads were normalized with DESeq2, and a pseudo count of 0.5 was added to enable plotting of zero values. Violin plots show the distribution across different TE groups: all TEs, TEs not significantly changed, TEs down-regulated, and TEs upregulated according to mRNA-seq in *Piwi6* KO over Ctrl cells. Data were analyzed using a Friedman test with Bonferroni-corrected pairwise Wilcoxon signed-rank post-hoc test on $\log_2$-transformed counts; ns, not significant; ** $P < 0.01$; **** $P < 0.0001$.

B-C) Volcano plots show the $\log_2$ fold changes in RNA polymerase II (RNA pol II) enrichment (B) and H3K9me3 enrichment (C) at TEs between *Piwi6* KO and WT cells. Vertical dashed lines highlight $\log_2 FC > 1$, and horizontal dashed line marks the threshold of statistical significance at adjusted *P* values ≤ 0.05. Individual elements are labeled according to the modified *Ae. Albopictus* repeat annotation [41].

S7

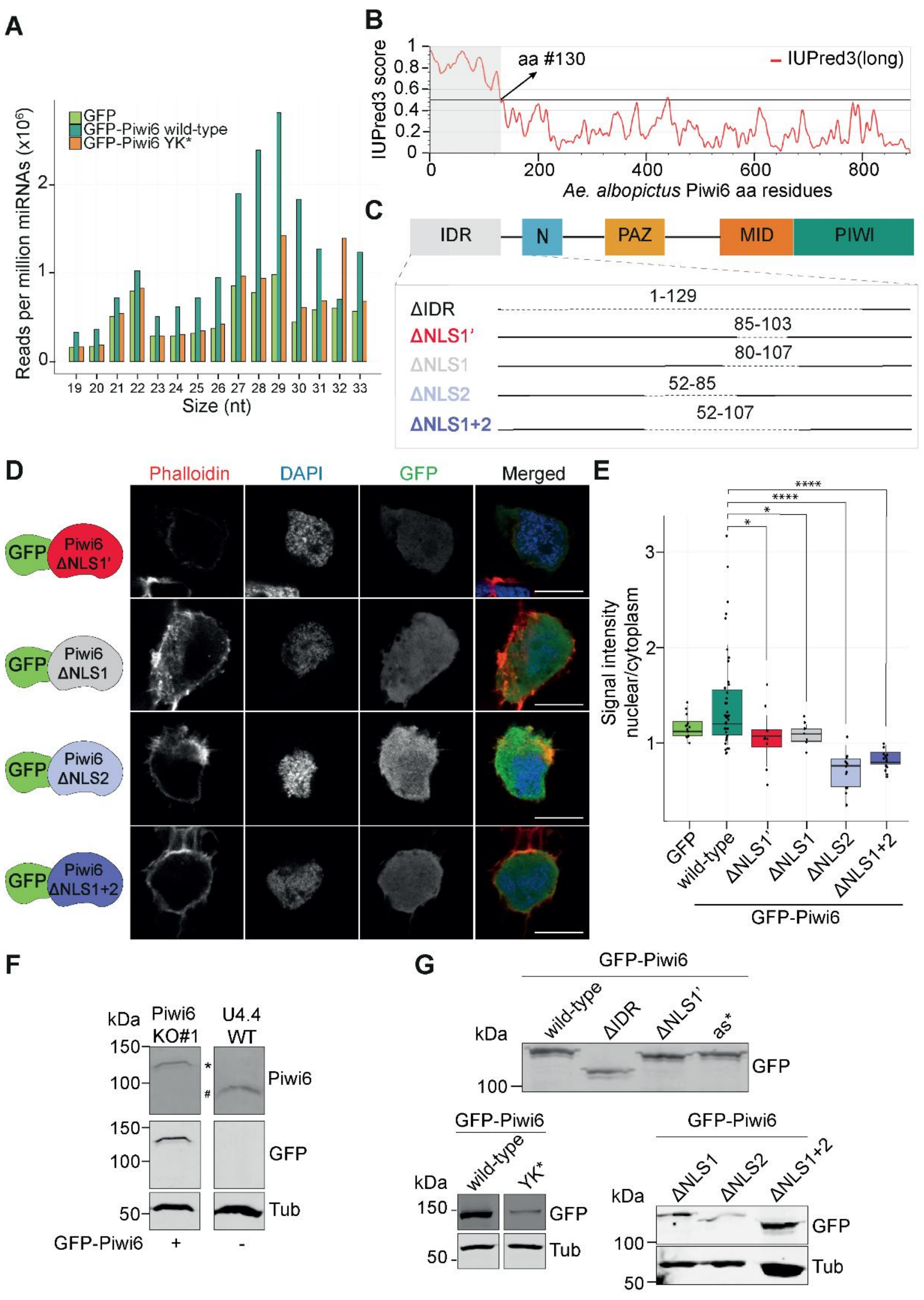

**Figure S7. Characterization of putative nuclear localization signals in Piwi6.**
A) Size profiles of small RNA reads mapping to the *Ae. albopictus* repeat annotation allowing 1 mismatch in libraries from immunoprecipitations of GFP, GFP-tagged wild-type Piwi6, or the GFP-tagged Piwi6-YK* mutant. Read counts were normalized to the total number of miRNAs (in millions).
B) Protein disorder plot of *Ae. albopictus* Piwi6 predicted by IUPred3 [37] using the default prediction (long disorder). Gray shading indicates the predicted IDR (aa residues 1–130).
C) Cartoon representation of mutations in the putative nuclear localization signals (NLSs) in the intrinsically disordered region (IDR) of Piwi6.
D) Representative confocal images showing the subcellular localization of GFP-tagged Piwi6 NLS-deletion constructs. DAPI and phalloidin staining were used to stain the nuclei and cytoplasm, respectively. Uncropped images are shown in Supplementary Figure S10. Scale bars represent 10 μm.
E) Quantification of the GFP signal intensity from panel C, presented as the nuclear to cytoplasm ratio. The centerline in boxplots represents the mean, box edges indicate the first and third quartiles, whiskers show maximum and minimum values. The values from GFP and GFP-tagged wild-type Piwi6 are the same as shown in 4D. Dots represent individual cells: GFP (n = 12), wild-type *Piwi6* (n = 41), ΔNLS1' (n = 12), ΔNLS1 (n = 7), ΔNLS2 (n = 13) and ΔNLS1+2 (n = 15). One-way ANOVA with Holm-Sidak's multiple comparisons post hoc test was performed to determine statistically significant differences with GFP-tagged wild-type Piwi6. ns; not significant; * $P < 0.05$; **** $P < 0.0001$. GFP-tagged wild-type Piwi6 and GFP samples are the same shown in Figure 5D.
F) Western blot analysis of Piwi6 in *Piwi6* KO cells transiently expressing a GFP-tagged Piwi6 transgene and non-transfected WT cells, detected by GFP and Piwi6 antibodies. The bands indicated with * and # correspond to endogenous Piwi6 and GFP-tagged Piwi6, respectively. Tubulin staining serves as a loading control. Individual panels were cropped from the same image, and the uncropped images are shown in Supplementary Figure S10.
G) Western blot analysis of Piwi6 in *Piwi6* KO cells transiently expressing the indicated GFP-tagged Piwi6 constructs stained with GFP antibody. Tubulin serves as a loading control. Images in the middle panel were cropped from the same image. Uncropped images are shown in Supplementary Figure S10.

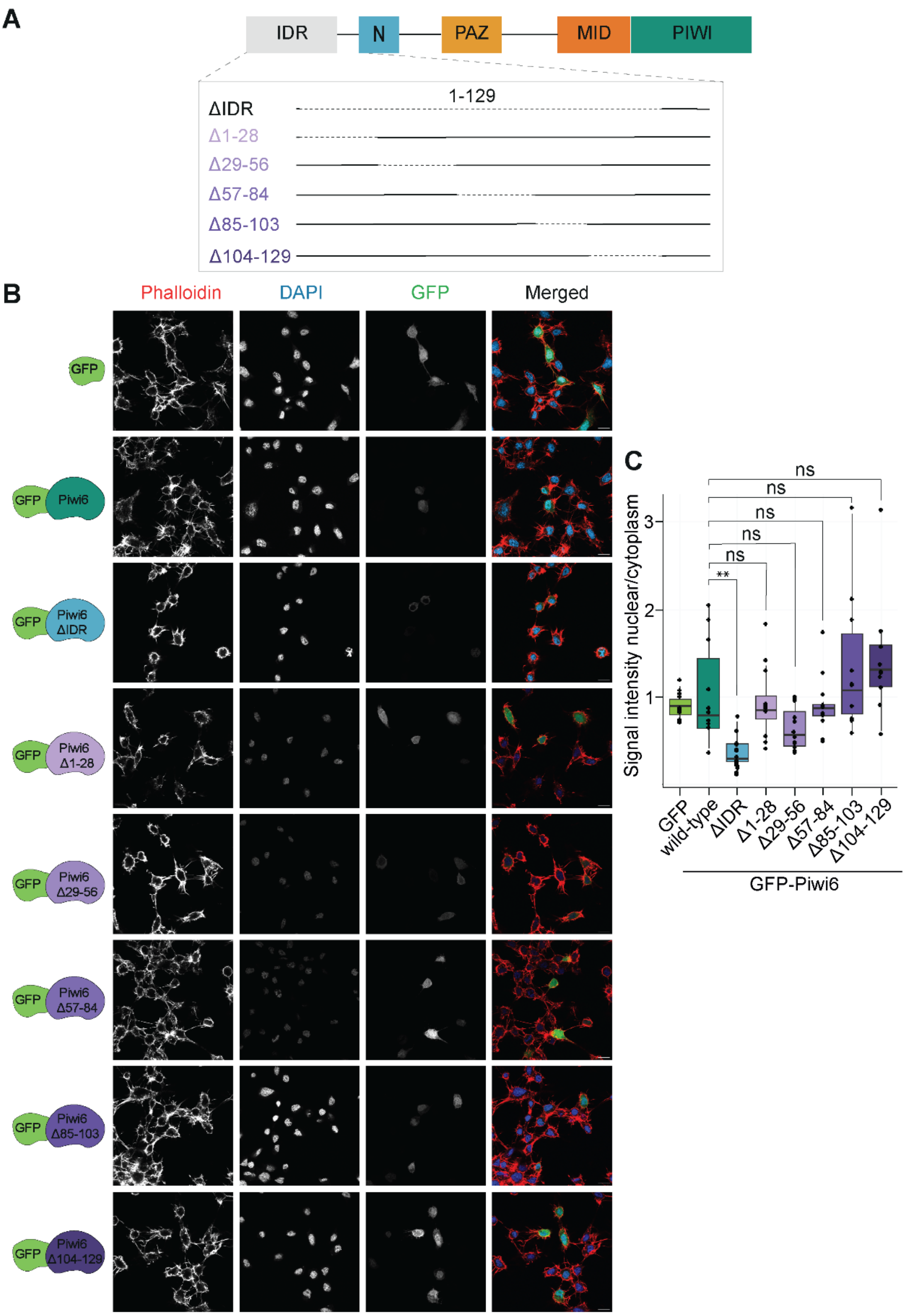
A
IDR
N
PAZ
MID
PIWI
1-129
ΔIDR
Δ1-28
Δ29-56
Δ57-84
Δ85-103
Δ104-129
B
Phalloidin
DAPI
GFP
Merged
GFP
Piwi6
Piwi6 ΔIDR
Piwi6 Δ1-28
Piwi6 Δ29-56
Piwi6 Δ57-84
Piwi6 Δ85-103
Piwi6 Δ104-129
C
Signal intensity nuclear/cytoplasm
ns
**
GFP
wild-type
ΔIDR
Δ1-28
Δ29-56
Δ57-84
Δ85-103
Δ104-129
GFP-Piwi6

**Figure S8. Subcellular localization of Piwi6 containing sequential deletions in the IDR**

A) Domain structure of *Ae. albopictus* Piwi6, showing deletion constructs within the IDR. Residues that are deleted are indicated with dashed lines.

B) Representative confocal images showing the subcellular localization of GFP, GFP-Piwi6, and the indicated mutants in *Piwi6* KO cells. DAPI and phalloidin staining were used to stain the nuclei and cytoplasm, respectively. Scale bars represent 10 µm.

C) Quantification of the GFP signal intensity from panel (B), presented as the ratio of nuclear to cytoplasmic expression. The centerline in the boxplots represents the mean, box edges indicate the first and third quartiles, whiskers show maximum and minimum values. Statistical significance was tested against GFP-tagged wild-type Piwi6. Dots represent individual cells: GFP (n = 10), wild-type Piwi6 (n = 11), ΔIDR (n = 14), Δ1-28 (n = 12), Δ29-56 (n = 14), Δ57-84 (n = 11), Δ85-103 (n = 10), Δ104-129 (n = 11). One-way ANOVA with Holm-Sidak's multiple comparisons post hoc test was performed to determine statistically significant differences with GFP-tagged wild-type Piwi6. ns; not significant; ** $P < 0.01$.

S9

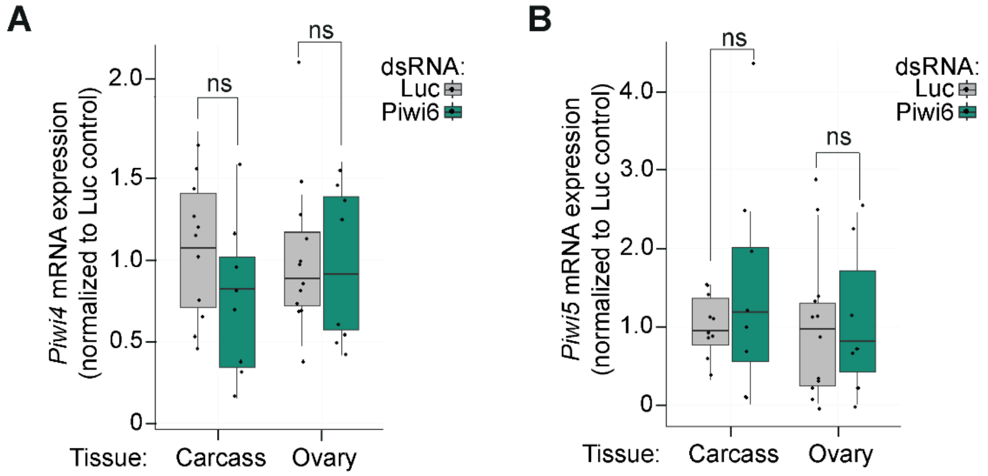


**Figure S9. Non-target gene expression upon *Piwi6* knockdown *in vivo***

A-B*Piwi4* (A) and *Piwi5* expression (B) in ovaries and carcasses from mosquitoes injected with dsRNA targeting *Piwi6* or firefly *luciferase* (Luc), assessed by RT-qPCR. mRNA expression of target genes was normalized to the expression of the housekeeping gene *RPL5* and expressed relative to the non-targeting control dsRNA (Luc). The centerline in the boxplots shows the mean, box edges represent the first and the third quartile, and the whiskers show maximum and minimum values. Differences in gene expression between Piwi6 and Luc dsRNA treated animals were not significantly different (two-tailed student's *t*-tests. Dots represent individual samples analyzed in carcass: Luc (n = 11) and Piwi4 (n = 8) and in ovaries: Luc (n = 12) and Piwi4 (n = 8) in A, and carcass: Luc (n = 10) and Piwi5 (n = 8) and in ovaries: Luc (n = 12) and Piwi5 (n = 7) in B.

**A.** Figure 1D uncropped

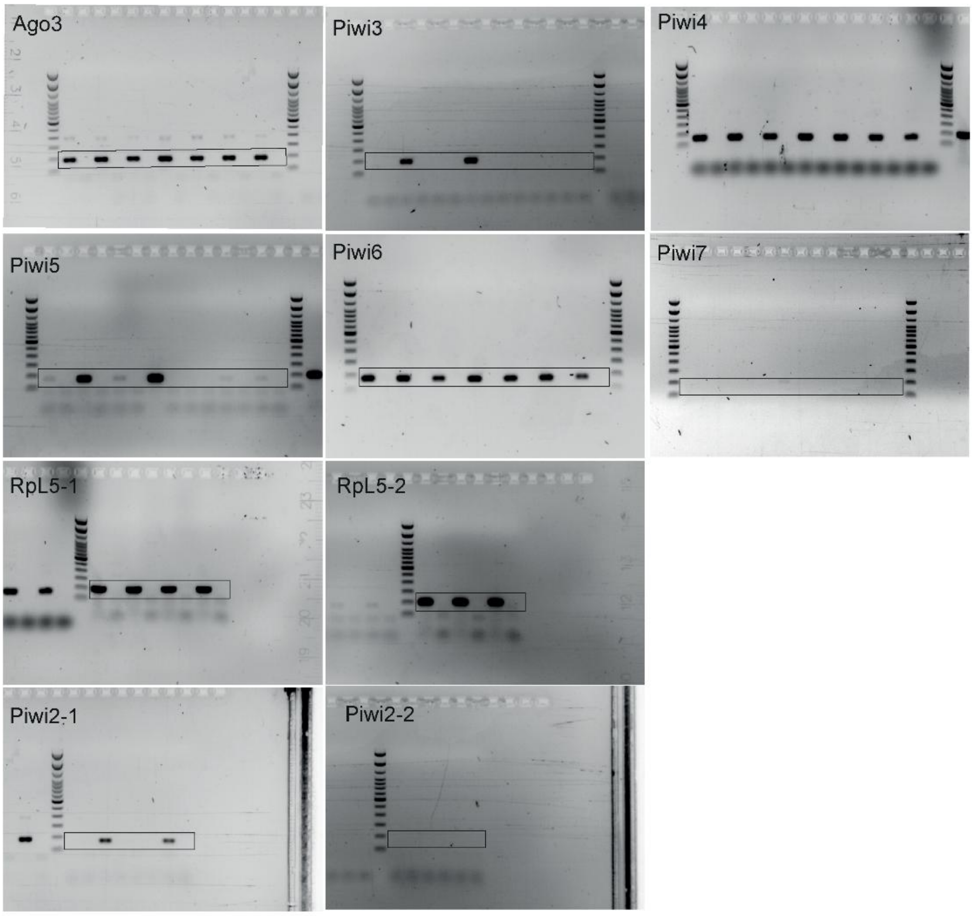


**B.** Figure 2A uncropped

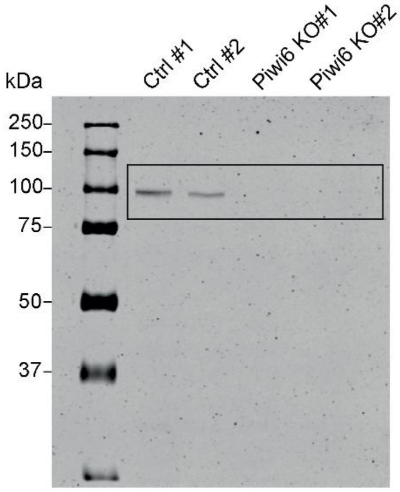

**C.** Figure 5B uncropped

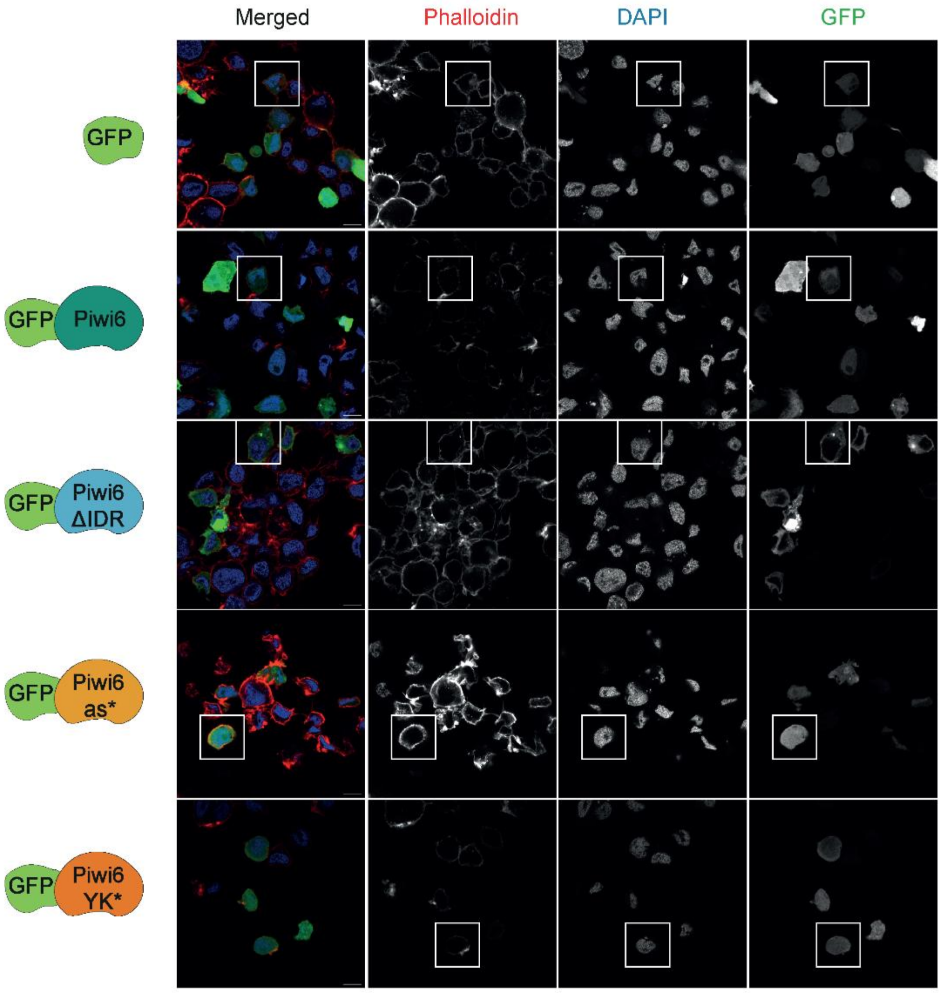


**D.** Figure 5D uncropped

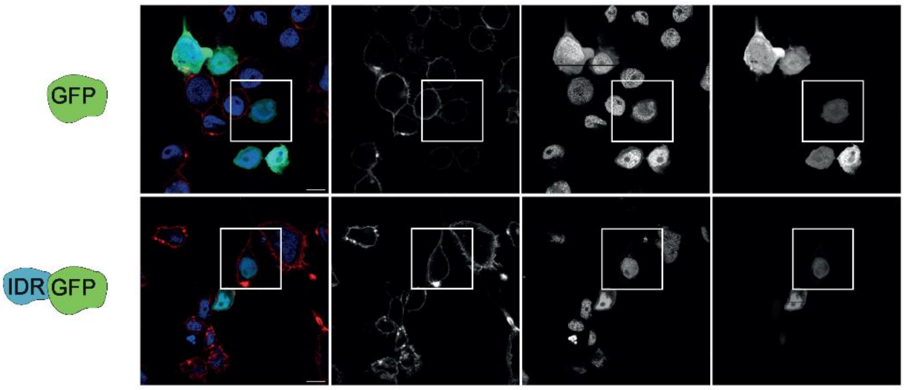

**E.** Supplementary Figure S7D uncropped

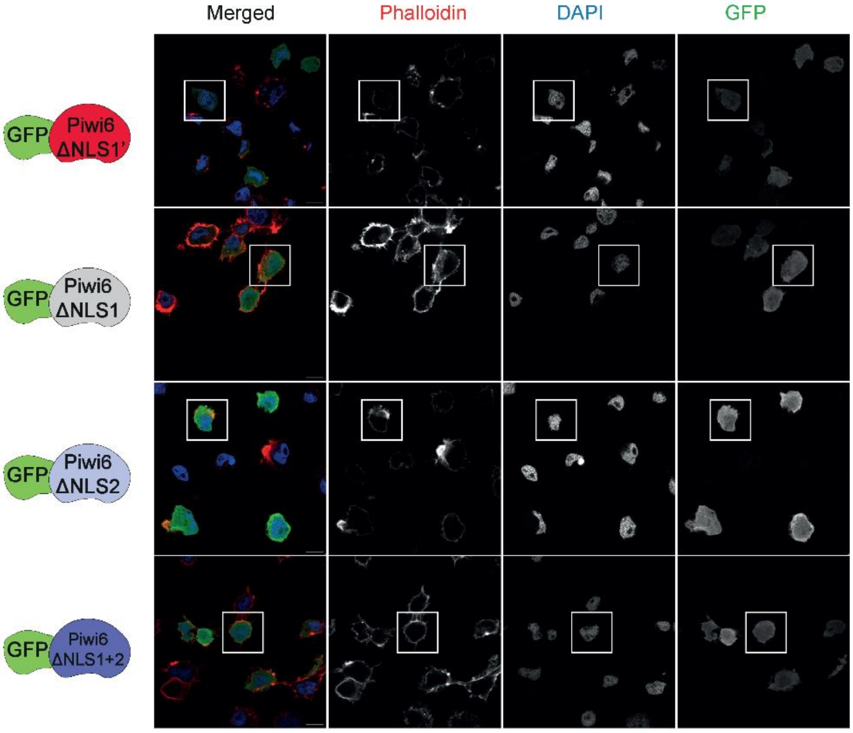


**F.** Supplementary Figure S7F uncropped

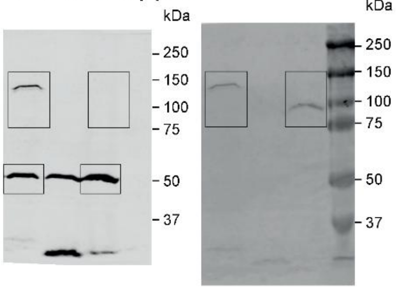


**G.** Supplementary Figure S7G uncropped

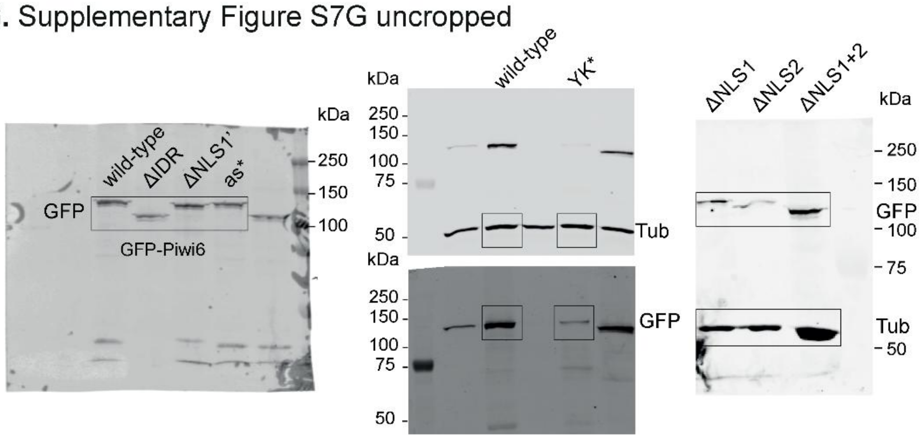

### Supplementary Tables

**Supplementary Table S1: Guide RNA sequences used for generating knockout lines**

| | |
|---|---|
| Aalb PIWI6 guide #1.1 | AATGATTAACCGTCACGCCATGGC |
| Aalb PIWI6 guide #1.2 | AACGCCATGGCGTGACGGTTAATC |

**Supplementary Table S2: Primers used for cloning**

| | |
|---|---|
| bb-pUb-GFP.AeP5_F | TAATAATAAAACCCAGCTTTCTTGT |
| bb-pUb-GFP.AeP5_R | GAAGCCTGCTTTTTGTACAAACT |
| Aalb.Piwi6-t1_Inf_F | AAAAAAGCAGGCTTCATGGCCGATAATCCCCAG |
| Aalb.Piwi6-t1_Inf_R | TGGGTTTTATTATTACTACAAAAAGTAAAGTTTCTTCTCC |
| Aalb.Piwi6-DelIDR-Inf_F | GCTTCATGCGACCGATCATGCTTCAGACC |
| Aalb.Piwi6-DelIDR-Inf_R | TCGGTCGCATGAAGCCTGCTTTTTGTACA |
| Aalb.Piwi6-DelNLS-Inf_F | AGAAGATGCGAGGAAACCGCTATTTGCC |
| Aalb.Piwi6-DelNLS-Inf_R | TTCCTCGCATCTTCTCCTGCTTAATTTCCG |
| Aalb.Piwi6-D656A-Inf_F | TGGATACGCTGTGTGCAAGGATTCCAAGG |
| Aalb.Piwi6-D656A-Inf_R | CACACAGCGTATCCAATAACCATCACCGAAGC |
| AlbP6-Δ80-107-Inf_F | CGGAAATTTATTTGCCGGAAATTGTGCATACCC |
| AlbP6-Δ80-107-Inf_R | GCAAATAAATTTCCGACTCCAATGCGG |
| AlbP6-Δ52-85-Inf_F | GTGATCACCAGGGCCGAGGTGAGCGTG |
| AlbP6-Δ52-85-Inf_R | GGCCCTGGTGATCACCTCTATATCCGCCTCCA |
| AlbP6-Δ52-107-Inf_F | GTGATCACTATTTGCCGGAAATTGTGCATACCC |
| AlbP6-Δ52-107-Inf_R | GCAAATAGTGATCACCTCTATATCCGCCTCC |
| AlbPiwi6-YK-Inf_F | TGCTTCGGCCATCGCAAAGAAGTGCTGCGTGGACC |
| AlbPiwi6-YK-Inf_R | GCGATGGCCGAAGCACGATCGGCCTTGTCGTTC |
| ins-pUb-IDRAlbP6GFP_F | GCTCGCATGCCCACCATGGCCGATAATCCCCAGG |
| ins-pUb-IDRAlbP6GFP_R | GCCCTTGCTCACCATTCCGGTGTTTCCTTGCTTGG |
| Aalb P6 Δ1-28 F | TACAACGGAGGATCTTCACG |
| Aalb P6 Δ1-28 R | AGATCCTCCGTTGTAGAAGCCTGCTTTTTGTAC |
| Aalb P6 Δ29-56 F | GATCGCCCGGAACATTCGC |
| Aalb P6 Δ29-56 R | AATGTTCCGGGCGATCGCCGCCACGCGATGAGGATC |
| Aalb P6 Δ57-84 F | CAGGGCCGAGGTGAGCGTG |
| Aalb P6 Δ57-84 R | CTCACCTCGGCCCTGGTAAGGCTTCTGCCGGTGATC |
| Aalb P6 Δ85-103 F | CGAGGAAACCGCTATTTGCC |
| Aalb P6 Δ85-103 R | ATAGCGGTTTCCTCGCATCTTCTCCTGCTTAATTTC |
| Aalb P6 Δ104-129 F | CCGATCATGCTTCAGACCAA |
| Aalb P6 Δ104-129 R | CTGAAGCATGATCGGCAGTCCACCACGACCACCC |

**Supplementary Table S3: Primers used in qPCR reactions**

| | |
|---|---|
| Alb_077280_qPCR_F | ATGCAGATGAACCCGATGAT |
| Alb_077280_qPCR_R | TCGACTTCTTTCCCTCGTTG |
| Alb_062793_qPCR_F | CCATGCAAGTTGAACCAATG |
| Alb_062793_qPCR_R | TCTCCGCAGAATTAGCAACA |
| Alb_049702_qPCR_F | GCAGTAGCAGCAGCAAAATG |
| Alb_049702_qPCR_R | TGTACTTTTCCAGGGCTTGC |
| Piwi6_qPCR_F | TCAACCCGGAGAGCACTTTC |
| Piwi6_qPCR_R | CCGCATGAGATGGAAATCCC |
| Piwi5_qPCR_F | GATCATCATCCGGTCAAGGC |
| Piwi5_qPCR_R | ATCTGCCGATCTTTGTTGTCC |
| AalRpL5-1_qPCR_F | TCGCTTACGCCCGCATTGAGGGTGAT |
| AalRpL5-1_qPCR_R | TCGCCGGTCACATCGGTACAGCCA |
| rnd6fam43LTR/Gypsy_qPCR_F | CACGGAGGGGAGTAGTTACG |
| rnd6fam43LTR/Gypsy_qPCR_R | CATGATGATTGCGTCTCAGG |
| SINV-NS4-1_qPCR_F | AACTCTGCCACAGATCAGCC |
| SINV-NS4-1_qPCR_R | GGGGCAGAAGGTTGCAGTAT |

**Supplementary Table S4: Primers used to generate templates for dsRNA production**

| | |
|---|---|
| Alb_077280-dsF-2 | GCCCGACGCCGAGGGTAACATTTCCCAGA |
| Alb_077280-dsR-2 | CGCCTCGGCTAGTGCAATCCTGCGTTCAG |
| Alb_062793-dsF-2 | GCCCGACGCTTCTTGTACCGTGCCCCTAC |
| Alb_062793-dsR-2 | CGCCTCGGCGCACAGGCATTGGGTTAGTT |
| Alb_049702-dsF | GCCCGACGCCCGCGTACATCAAGAAAGAA |
| Alb_049702-dsR | CGCCTCGGCCAGTGGATTTCTCGGCTTTT |
| Aalb.Piwi6_dsT7_F | GCCCGACGCACGATTCACCCGTCGTCTAC |
| Aalb.Piwi6_dsT7_R | CGCCTCGGCATGCTGGTTTACGGTCGAGT |

**Supplementary Table S5: Primers used in PCR reactions**

| | |
|---|---|
| Ago3_qPCR_F | CGAATGTTCCGCATCGATGG |
| Ago3_qPCR_R | AGAATGTCCGGCGAATGGTT |
| Piwi2_qPCR_F | GAAGCGATTTTCGAATGATGCG |
| Piwi2_qPCR_R | AGACGCTTGTCCAACTCCATC |
| Piwi3_qPCR_F | CCGACGGAGAAAGGTGTCC |
| Piwi3_qPCR_R | TGGATTTCGAACCGCATTCC |
| Piwi4_qPCR_F | GCTAATGGCGTACAAGCTGG |
| Piwi4_qPCR_R | CTGAATCATGCAGCCCAGGA |
| Piwi5_qPCR_F | GATCATCATCCGGTCAAGGC |
| Piwi5_qPCR_R | ATCTGCCGATCTTTGTTGTCC |
| Piwi6_qPCR_F | TCAACCCGGAGAGCACTTTC |
| Piwi6_qPCR_R | CCGCATGAGATGGAAATCCC |
| Piwi7_qPCR_F | GCGGCGGAATTTCAATTTAATGC |
| Piwi7_qPCR_R | ATCGTTTCCGGTTGAAGCAC |

**Supplementary Table S6: Primers used in PRO-seq library preparation**

| | |
|---|---|
| 3' RNA adapter | [Phos]GAUCACGGAUCGUCGGACUGUAGAACUCUGAAC[InvdT] |
| 5'RNA adapter | [InvdT][dC][dC][dT][dT][dG][dG][dC][dA][dC][dC][dC][dG][dA][dG][dA][dA][dT][dT][dC][dC][dA]NNNNNNC |
| PROseq _RP1 | AATGATACGGCGACCACCGAGATCTACACGTTCAGAGTTCTACAGTCCGA |
| PROseq_RPI-2 | CAAGCAGAAGACGGCATACGAGATACATCGGTGACTGGAGTTCCTTGGCACCCGAGAATTCCA |
| PROseq_RPI-3 | CAAGCAGAAGACGGCATACGAGATGCCTAAGTGACTGGAGTTCCTTGGCACCCGAGAATTCCA |
| PROseq_RPI-4 | CAAGCAGAAGACGGCATACGAGATTGGTCAGTGACTGGAGTTCCTTGGCACCCGAGAATTCCA |
| PROseq _RPI-10 | CAAGCAGAAGACGGCATACGAGATAAGCTAGTGACTGGAGTTCCTTGGCACCCGAGAATTCCA |
| PROseq _RPI-11 | CAAGCAGAAGACGGCATACGAGATGTAGCCGTGACTGGAGTTCCTTGGCACCCGAGAATTCCA |
| PROseq _RPI-12 | CAAGCAGAAGACGGCATACGAGATTACAAGGTGACTGGAGTTCCTTGGCACCCGAGAATTCCA |

**Supplementary Table S7: Primers used in CUT&Tag library preparation**

| | |
|---|---|
| CUT&Tag_i5-S501 | AATGATACGGCGACCACCGAGATCTACACTAGATCGCTCGTCGGCAGCGTCAGATGTGTAT |
| CUT&Tag_i5-S502 | AATGATACGGCGACCACCGAGATCTACACCTCTCTATTCGTCGGCAGCGTCAGATGTGTAT |
| CUT&Tag_i7-N701 | CAAGCAGAAGACGGCATACGAGATTCGCCTTAGTCTCGTGGGCTCGGAGATGTG |
| CUT&Tag_i7-N702 | CAAGCAGAAGACGGCATACGAGATCTAGTACGGTCTCGTGGGCTCGGAGATGTG |
| CUT&Tag_i7-N703 | CAAGCAGAAGACGGCATACGAGATTTCTGCCTGTCTCGTGGGCTCGGAGATGTG |
| CUT&Tag_i7-N704 | CAAGCAGAAGACGGCATACGAGATGCTCAGGAGTCTCGTGGGCTCGGAGATGTG |
| CUT&Tag_i7-N705 | CAAGCAGAAGACGGCATACGAGATAGGAGTCCGTCTCGTGGGCTCGGAGATGTG |
| CUT&Tag_i7-N706 | CAAGCAGAAGACGGCATACGAGATCATGCCTAGTCTCGTGGGCTCGGAGATGTG |

**Supplementary Table S8: *AalERV1* elements in *Ae. albopictus* genome (0-based)**

| contig name | start site | end site | # of ORFs | orientation |
|---|---|---|---|---|
| NW_021839020.1 | 49440331 | 49447338 | 5 | + |
| NW_021838549.1 | 195388 | 202477 | 6 | + |
| NW_021838443.1 | 3566184 | 3573318 | 4 | + |
| NW_021838465.1 | 97014947 | 97022302 | 3 | - |
| NW_021838190.1 | 1015081 | 1022603 | 7 | + |
| NW_021837600.1 | 22350691 | 22358291 | 3 | + |
| NW_021837156.1 | 35936832 | 35944534 | 3 | - |
| NW_021837045.1 | 92943148 | 92950889 | 11 | - |
| NW_021837045.1 | 146399654 | 146407556 | 3 | + |
| NW_021837046.1 | 39660383 | 39668274 | 5 | + |
| NW_021839020.1 | 15603126 | 15611018 | 6 | + |
| NW_021837045.1 | 94945995 | 94953890 | 5 | + |
| NW_021838909.1 | 8962227 | 8970122 | 5 | - |
| NW_021839020.1 | 34709752 | 34717647 | 7 | - |
| NW_021837489.1 | 25242329 | 25250225 | 3 | - |
| NW_021837489.1 | 42924452 | 42932341 | 4 | + |
| NW_021838576.1 | 97882906 | 97890807 | 3 | - |
| NW_021838909.1 | 1829801 | 1837701 | 3 | - |
| NW_021837046.1 | 21962233 | 21970135 | 3 | + |
| NW_021838153.1 | 7827714 | 7835618 | 3 | + |
| NW_021838762.1 | 317462 | 325360 | 3 | + |
| NW_021837069.1 | 612770 | 620667 | 4 | - |
| NW_021838576.1 | 14281325 | 14289216 | 9 | - |
| NW_021839121.1 | 154880 | 162779 | 6 | + |
| NW_021839086.1 | 2012655 | 2020555 | 4 | - |
| NW_021838868.1 | 13869 | 21777 | 5 | + |
| NW_021837045.1 | 64008423 | 64016326 | 6 | + |
| NW_021837600.1 | 5396367 | 5404268 | 7 | + |

**Supplementary Table S9: ORF sequences of NW_021838153.1:7827714-7835618**

>ORF_906-2387_493aa
MARYDGNRDLLAAENEALQAQLEQAAADMVEQNRQVIELQHQLAAGQDVQNQLDNALLQINQKDQQ
IANLRAATLNNFSDVESILKSLQTPQILRDFPCFTGDPVKLHSFIKSIDKLMPTLERAKGTPAFDVWMHA
IRSKITHDADAALELYGTETNWDDIKNTLITHFSDKRDEISLTRDLFKLKQGDSVQDFYRDVSHSISLLIN
LLQVNEKVSVIIAAKKQFYQDLGLKVFLSGLKEPLGPIIRAQAPKTLKEALRLCLEEDNYNYVKNSFQSIS
KTSNPISLPPKPMQKSFQFPPPRPFNFQKPPQYQKPFQFQSTPRPQFNNGPRPFQFNTAQKPFQFR
SFQQNQFQRPFQYQRNPFMPNTFQRNFPKPQPKPTPMEVDPSLSKQVNYMNRPHYQVEEFDECFD
YPNYSEPYYYPEYTEYSPYPQYSEYPEYLDYSEYFDEDPNQSQVHNNNINLPKVTLPAIKECEQNNVQ
VDDINFRERTEDGHPT

>ORF_2384-5806_1140aa
MNNFIPYIEVDTTKGPLKFLIDTGANKNYISPKHVNQAKFTDPLTVHNINGVHTINQSVHFNPFYKIANT
QKLQFYVFDFHPFFDGLIGYEALQDLKANIITSTNELQLPSGNIKMLRKYPCSTQVQLNANETKFIHFPV
NFEKGDFFVSHDTQIHDSIFVHAGLYKAENNHAFLAITNHRNNPIQIEIFKPFVSELNNFESQSYPHPVR
PINRQLFNQLRLDHLNSEEKPKLLKLIADNQDVFYLPGESLTFTNAIKHSIKTKDEMPIYCKSYRYPFCH
KDEVQKQINEMLEQGIIRHSISPWTSPVWIVAKKMDASGLKKWRLVIDYRKLNDKTIDDRYPIPNITDILD
KLGRCNYFTTLDLASGFHQIEVEESDIPKTAFNVENGHYEFLRMPFGLKNAPSTFQRVMDNILREHIGV
RCFIYMDDIIVFSTSLQEHLDNLRKIFNTLSKYNMKVQLDKCEFLRKEVAFLGHIVSTDGVKPNPEKISIIK
NWPLPKNERELRGFLGILGYYRKFIKDFAKIAKPLTNSLRKGETIIHSENFVKAFQKCKSILTSSDILQYP
DFDKSFILTTDASNFAIGAVLSQGPLGKDKPIAFASRTLNKTEENYSTIEKELLAIVWACKHFRPYLYGR
KFILYTDHKPLTYSLNLKEPNQRLIHWRLTLSEFDYEIRYKPGKQNVVADSLSRIFQEVNINEDETSSSN
DTMHSADTDDSELIPCTEIPINYFSNQIILKIGNEESNIYEEVFPRVYRRTITKFIFGVPFIIKIFKEYMDTRR
TNCILCPENVINIIQFVYKNYFSRCRTFKVKISQTMRIDVRTEVEQDTLIENTHETSHRGILENLCEIRNRF
FFPKMKNKIRKYIILCEICNKMKYERKPYKIKLGETPIPKQPLDIIHIDIFISQPNLFLSIVDKLSRFGFLIPIK
SRTTIDIRKGLIKFISQFGSPKLIVSDNEPAFKSIEIRDLLNNLNIQQYFTPTNYSEVNGIVERFHSTLSEIY
RCNKHKYENLSSKEMFLIACSLYNNAIHTSTKLKPREVFYGIKDGEERPLDRAQLIEYRDKIYDETITKIT
STQQEQHKNRNASREDPPALEPDQQAFNRRQGIKAKTKEWFEKVKVTSDRNQTYKDDKNRKLHKTK
MRRIRK

>ORF_5846-7612_588aa
METTTLLLFLISINCMLQPIHCTIQIMDLNQNPGLLALTAGESFMKVGEHRIYHTIDFDLYEPTFTKLKLIT
KELRNYGNFTELNDVLSTKLFNLKNLYFGLQMKTRQKRGLMDFVGTGIKFLTGNMDHQDYIDISRDLD
DLTTKSNQLIRENNEQRKINYDMQDRINRLIKQINNQQSLIMNAINNNQSGSLQKQIKILESIINVNIQLDH
LISHFKDISESIHLAKVNIISKHILHPDELSFSIDRLEEKGIVIHNFEQVYDFLEISAFYNMTKLVFVVKIPSL
KNHTYSRLILEALPIENKILNLPASTAMISNNETYFITKECRNIEEYRICNQKDLLDLSNDTCFTQLLRGM
TGKCTFKMYNQQKEFKQITNNHIIVKSKSPVNIRSNCEITNRNLTGSYLIEFRNCSIILDSTKFENTEIFKT
EMPLIVPLDGLHIEKQIIESSIEDLQIQNRNHLETLSTSHQIHHYSSISLSVFSILGIITLIILYKVKTKNVNINF
RRKPFLEEVSNNQAKDESLELREISSSPRIQSTLLSTRDVSTSRRGVVTSNPFPLTTIKPLPPIMPIFVNK
PLSLSKPETQSS

**Supplementary Table S10: Gene accession numbers used in Phylogeny Analysis**

| | ***Ae. aegypti*** | ***Ae. albopictus*** | |
|---|---|---|---|
| **Gene name** | **VectorBase ID** | **LOC number** | **VectorBase ID** |
| *Ago3* | *AAEL007823* | *LOC109433074*<br>*LOC109411297* | *Ago3a: AALFPA_060827*<br>*Ago3b: AALFPA_062933* |
| *Piwi3* | *AAEL013692* | *LOC109401827* | *AALFPA_073526* |
| *Piwi2* | *AAEL008098* | *LOC109401826* | *AALFPA_061874* |
| *Piwi4* | *AAEL007698* | *LOC109404625* | *AALFPA_053057* |
| *Piwi5* | *AAEL013233* | *LOC109413120* | *AALFPA_066630* |
| *Piwi6* | *AAEL013227* | *LOC109418558* | *AALFPA_065618* |
| *Piwi7* | *AAEL006287* | *LOC109422824*<br>*LOC115260637* | *Piwi7a: AALFPA_073388*<br>*Piwi7a:AALFPA_050058* |
| **Gene name** | **FlyBase ID** | | |
| *Dm. Piwi* | CG6122 | | |
| *Dm. Aub* | CG6137 | | |
| *Dm. Ago3* | CG40300 | | |
| **Gene name** | **Ensembl Metazoa ID** | | |
| *Bm. Ago3* | GeneID_100125337 | | |
| *Bm. Siwi* | GeneID_100125336 | | |

**Supplementary Table S11:** ***AalERV1*** **insertions in piRNA clusters***

| Cluster coordinates (target) | percentage nt identity | alignment length | expect value |
|---|---|---|---|
| NW_021837046.1: 21962257-21970095 (U4.4cluster96) | 97.847 | 7897 | 0.0 |
| NW_021838042.1: 25491379-25512840 (U4.4cluster53) | 97.627 | 3203 | 0.0 |
| NW_021838042.1: 25491379-25512840 (U4.4cluster53) | 96.784 | 1710 | 0.0 |
| NW_021838042.1: 25491379-25512840 (U4.4cluster53) | 94.899 | 1784 | 0.0 |
| NW_021838042.1: 25491379-25512840 (U4.4cluster53) | 94.038 | 1258 | 0.0 |
| NW_021838153.1: 164445001-164574032 (U4.4cluster2) | 88.989 | 1326 | 0.0 |
| NW_021837998.1: 285258-339963 (U4.4cluster18) | 88.536 | 1352 | 0.0 |

*Clusters are defined in Qu, Betting et al. 2023. doi: 10.1016/j.celrep.2023.112257.

**Supplementary Table S12: CUT&Tag H3K9me3 data**

The columns contain the following information: A, *Aedes albopictus* repeat annotations; B, Mean of normalized counts for all samples; C, $Log_2$ fold change (MLE): condition KO vs WT ; D, Standard error: condition KO vs WT; E, Wald statistic: condition KO vs WT; F, Wald test *P*-value: condition KO vs WT; G, Benjamini-Hochberg adjusted *P*-values.

**Supplementary Table S13: CUT&Tag RNA polymerase II data.**

The columns contain the following information: A, *Aedes albopictus* repeat annotations; B, Mean of normalized counts for all samples; C, $Log_2$ fold change (MLE): condition KO vs WT ; D, Standard error: condition KO vs WT; E, Wald statistic: condition KO vs WT; F, Wald test *P*-value: condition KO vs WT; G, Benjamini-Hochberg adjusted *P*-values.

**Supplementary Table S14: Small RNA read counts mapped to *Aedes albopictus* repeat annotations.**

The columns contain the following information: A, *Aedes albopictus* repeat annotations; B,C,D, Raw read counts of libraries from immunoprecipitations (IPs) of GFP, GFP-tagged wild-type Piwi6 transgene, and the GFP-tagged Piwi6-YK* mutant, respectively; E,F,G, Normalized counts of libraries from IPs of GFP, GFP-Piwi6, and GFP-Piwi6-YK* mutant, respectively; H; ratio of normalized counts in libraries from GFP-Piwi6 IP over GFP IP.
